# Cryo-ET of HIV reveals Env positioning on Gag lattice and structural variation among Env trimers

**DOI:** 10.1101/2021.08.31.458345

**Authors:** Vidya Mangala Prasad, Daniel P. Leaman, Klaus N. Lovendahl, Jacob T. Croft, Mark A. Benhaim, Edgar A. Hodge, Michael B. Zwick, Kelly K. Lee

## Abstract

HIV-1 Env mediates viral entry into host cells and is the sole target for neutralizing antibodies. However, Env structure and organization in its native virion context has eluded detailed characterization. Here we used cryo-electron tomography to analyze Env in mature and immature HIV-1 particles. Immature particles showed distinct Env positioning relative to the underlying Gag lattice, providing insights into long-standing questions about Env incorporation. A 9.1Å sub-tomogram averaged reconstruction of virion-bound Env in conjunction with structural mass spectrometry revealed unexpected features, including a variable central core of the gp41 subunit, heterogeneous glycosylation between protomers plus a flexible stalk that allows Env tilting and variable exposure of neutralizing epitopes. Together, our results provide an integrative understanding of HIV assembly and structural variation in Env antigen presentation.

## Introduction

Human Immunodeficiency Virus-1 (HIV-1) continues to infect nearly two million people worldwide each year, with no vaccine available against the virus (Pandey and Galvani 2019). The envelope glycoprotein (Env) on the surface of HIV-1 is an essential viral entry machine that mediates binding to host cell receptors and subsequent membrane fusion. As the sole target for neutralizing antibodies, Env is also of singular importance for vaccine design efforts (van Gils and Sanders 2013). Env is translated as a precursor gp160 protein, which trimerizes, and is cleaved into gp120 and gp41 subunits that are primarily responsible for receptor binding and fusion, respectively. The gp120 and gp41 subunits remain as non-covalently associated heterodimers with gp41 embedded in the viral membrane via its transmembrane domain (TMD) and the C-terminal cytoplasmic domain (CTD) in the viral lumen. Upon incorporation into budding virions, Env CTD interacts with the matrix domain (MA) of immature Gag polyprotein which assembles as a membrane-associated lattice on the inner side of the viral bilayer (Checkley, Luttge et al. 2011, Tedbury and Freed 2014).

Recent insights into Env structure have primarily come from studies of engineered, truncated forms of the trimer ectodomain (such as SOSIPs) (Ward and Wilson 2015, Stewart-Jones, Soto et al. 2016), while details of Env structure in its viral membrane context, including its organization relative to other viral components, has remained elusive. In order to understand how Env is assembled on virions and how it differs structurally from soluble trimers, we carried out cryo-electron tomography (cryo-ET) with sub-tomogram averaging, and structural mass spectrometry analysis of Env displayed on immature and mature Gag-bearing HIV-1 virus-like particles. These studies advance our knowledge of membrane-bound Env via insights gained from cryo-ET of intact viral particles.

## Results and Discussion

Virion particles bearing ADA.CM Env, a variant of subtype B isolate ADA that was selected for stability (Leaman and Zwick 2013), were produced in human HEK 293T cells as previously described (Stano, Leaman et al. 2017) (Figure 1A, B). These membrane enveloped virus-like particles, referred to as high Env (h)VLPs, display elevated levels of Env trimers that are fully functional for mediating receptor binding and entry. Env on hVLPs retain the antigenic profile of full-length Env (Figure S1 and Table S1), despite a C-terminal truncation of 102 amino acids in the CTD (Stano, Leaman et al. 2017). hVLPs assemble around a functional Gag layer that undergoes proteolytic maturation and have the composition of authentic HIV-1 virions but are replication incompetent because the packaged genome lacks the Env gene (Leaman and Zwick 2013, Stano, Leaman et al. 2017). Purified hVLPs were additionally treated with aldrithiol-2 (AT-2), a mild oxidizing agent that interrupts nucleocapsid interaction with RNA, to ensure non-infectivity (Rossio, Esser et al. 1998). Antibody binding experiments showed that the antigenicity of hVLP-Env was unaffected by AT-2 treatment (Figure S2). Despite the lack of infectivity of AT-2 treated hVLPs, they induced syncytia formation and cytotoxicity in cells, similar to non-AT-2 treated hVLPs, in accordance with a previous report (Rossio, Esser et al. 1998), showing that AT-2 treated HIV-1 virions bear functional Env that are capable of cell entry (Figure S2). These results indicate that the Env conformation on the hVLPs was not significantly altered by AT-2 as the antigenicity and viral entry properties of Env are retained. The high density of surface Env on hVLPs allowed us to gather sufficient cryo-ET data to obtain a 9.1Å resolution structure of Env in a membrane context on these virus-like particles (Figure 1C and S3).

**Figure 1.**
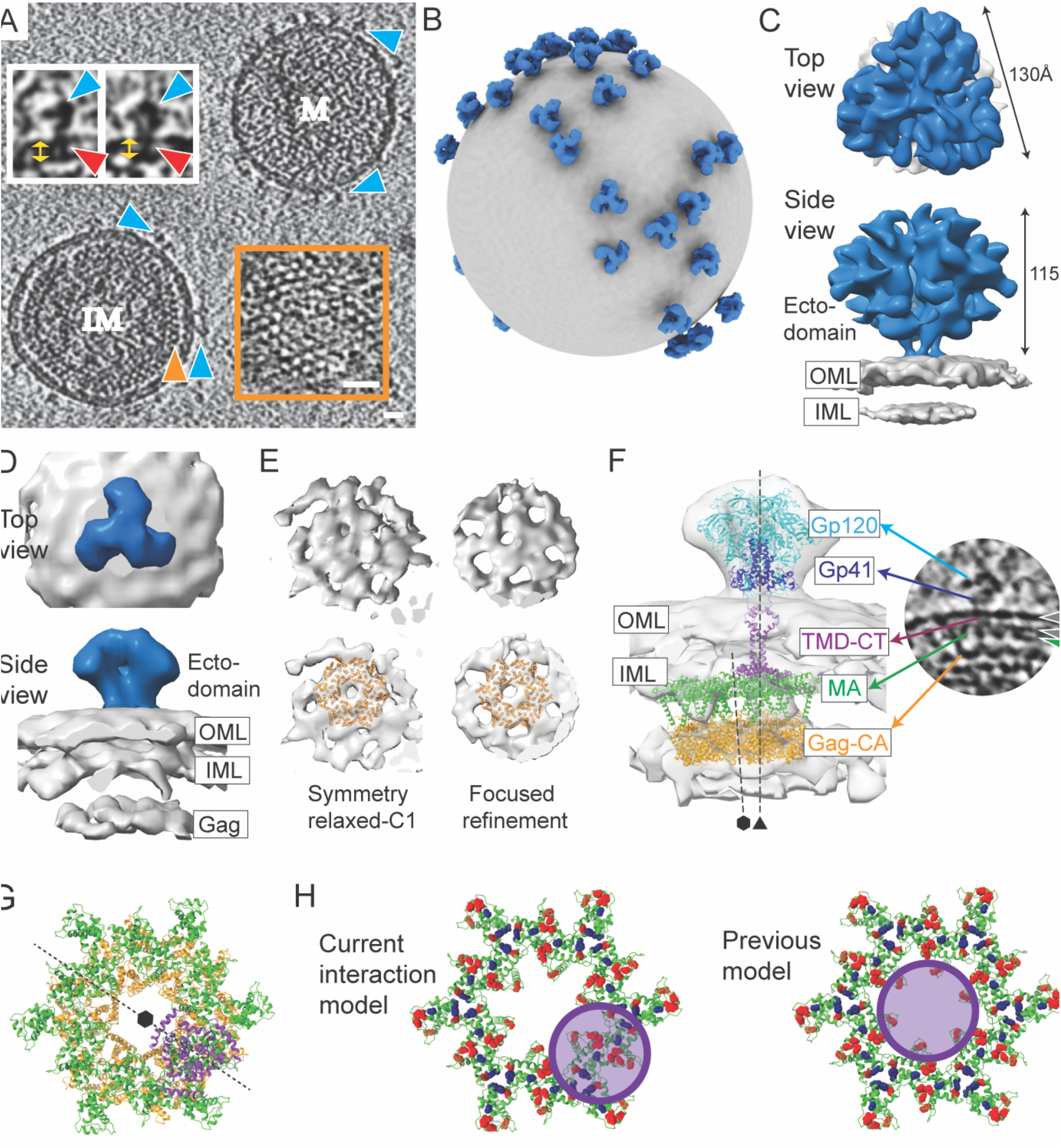
Structural analysis of hVLP-Env. **(A)** Tomogram slice showing a mature (M) and immature hVLP (IM). Blue arrows indicate Env. Orange arrow indicates Gag layer in IM with zoomed inset of Gag top view on bottom right. Scale bars indicate 200 Å. Top left inset shows close-up of Env with red arrows denoting TMD and yellow arrows denoting membrane bilayer. (For antigenic profile of hVLPs, see Figures S1 and S2). (**B)** Representative model of mature hVLP with averaged Env structure (in blue) placed onto original particle coordinates. Membrane surface is shown in grey. (**C)** Sub-tomogram averaged Env structure at 9.1Å resolution (for resolution related statistics, see Figure S3). OML and IML indicate outer and inner membrane leaflets, respectively. **(D)** Sub-tomogram averaged Env structure from immature VLPs. OML and IML indicate outer and inner membrane leaflets, respectively. (**E)** Top views of immature Gag layer from relaxed C1 symmetry map (top left) and after focused refinement (top right) (see Figure S4 for details). Bottom panels show the same with fitted atomic structure of hexameric Gag-CA (PDB: 4USN). (**F)** Left: Composite model of Env-Gag structures in immature particles. Atomic coordinates of Env ectodomain with Gp120 (cyan) and Gp41 (dark blue), TMD and CTD in purple (PDB: 6UJV), Gag-MA hexamer (PDB: 1HIW) in green and Gag-CA hexamer in orange are shown. Right: Tomogram slice of Env from an immature particle with protein components labeled. Grey and green arrows indicate membrane bilayer and Gag-MA respectively. (**G)** Top view of CTD (purple) and Gag-MA (green) interface. CTD lies on the rim of underlying Gag-CA hexamer (orange). **(H)** Model of Env-CTD interaction with Gag-MA. Ribbon diagram of Gag-MA hexamer (green) (PDB ID: 1HIW) (Hill, Worthylake et al. 1996) is shown. Residues that reduce Env incorporation when mutated (13L, 17E, 31L, 35V, 99E and 75LG) are highlighted as red spheres (Alfadhli, Staubus et al. 2019). Residues that reportedly suppress Env incorporation defects (62QR, 35VI and 44FL) are shown as blue spheres (Alfadhli, Staubus et al. 2019). Position and area covered by Env CTD structure (based on PDB ID: 6UJV (Piai, Fu et al. 2020)) in our current model (left) and previously predicted models (right) is shown as purple circle. In previous models (right panel), Env CTD position was predicted to be at the center of the Gag-MA hexamer. In this configuration, Env CTD directly interacts with only a few residues that line the inner circumference of Gag-MA hexamer. Thus, the effect of the majority of involved residues on Env incorporation would be allosteric. In the current model (left panel) based on our cryo-ET data, the Env CTD makes direct contacts with all Gag-MA residues determined to affect Env incorporation.

### Env structure from immature particles reveal interactions with Gag

Though the hVLPs in our sample population were predominantly mature, approximately 3% of the particles appeared immature, exhibiting a clear, assembled Gag lattice beneath the viral membrane (Figure 1A). Env sub-tomogram volumes from only these immature particles were averaged together to give density maps at ∼31 Å (C3 symmetry) and ∼34 Å (C1 symmetry) (Figure 1D and S4). The Env structure from immature particles closely resembles the sub-nanometer Env structure generated from the total hVLP population (Figure 1C) and other reported Env ectodomain structures (Kwon, Pancera et al. 2015, Ward and Wilson 2015). In the averaged immature Env map, a third density layer is seen underneath the membrane bilayer corresponding to the Gag lattice position (Figure 1A, D and Figure S4B, C). Focused refinement of this internal third layer revealed a distinct structure that clearly resembles a lattice formed by the capsid domain (CA) of immature Gag polyprotein (Figure 1E and Figure S4D) (Schur, Hagen et al. 2015). Thus, structures of trimeric Env ectodomain (Pan, Peng et al. 2020) and hexameric Gag-CA (Schur, Hagen et al. 2015) were fitted as rigid bodies into corresponding densities in the averaged map (Figure 1F). No clear density for the Env TMD-CTD was observed in the averaged structure, even though the TMD is seen in individual raw tomograms (Figure 1F), suggesting that the TMD is not strictly aligned with the ectodomain or Gag-CA. The structure of the TMD-CTD (PDB ID: 6UJV) was thus placed into the map based on its position relative to the Env ectodomain (Figure 1F) (Piai, Fu et al. 2020).

The N-terminal MA domain of immature Gag polyprotein assembles along the inner membrane leaflet as a hexameric lattice of trimers (Frank, Narayan et al. 2015, Tedbury, Novikova et al. 2016, Qu, Ke et al. 2021) and is connected by a flexible linker to the Gag-CA lattice underneath (Tedbury and Freed 2014). Regularly spaced puncta corresponding to the Gag-MA domain can be discerned directly underneath the inner leaflet and Env ectodomains in immature hVLP tomograms (Figure 1F and 2A). However, distinguishable density for the Gag-MA layer was not seen in the averaged immature map implying that the Gag-MA lattice (Tedbury and Freed 2014, Frank, Narayan et al. 2015, Tedbury, Novikova et al. 2016) may have subtle variations in arrangement that deviate from Gag-CA or Env symmetry axes. Hence, based on the expected position of Gag-MA relative to the Gag-CA lattice (Frank, Narayan et al. 2015, Tedbury, Novikova et al. 2016), the MA lattice structure was placed into the map (Figure 1F). The resulting juxtaposition of Env CTD and Gag-MA occurs right at the bottom of the inner membrane as would be expected from the raw tomograms (Figures 1F and 2). From this composite model, the relative position of Env CTD, when viewed normal to the membrane, is on the rim of the hexameric Gag-CA lattice along its 2-fold axis (Figure 1G). This observation can also be corroborated in the raw tomograms (Figure 2A and B). Thus, the trimeric Env CTD does not lie in the central hole formed by the immature Gag lattice as has been hypothesized (Tedbury and Freed 2014, Tedbury, Novikova et al. 2016), but instead is positioned on its hexameric rim (Figure 1G, H). In this configuration, the modeled Env CTD makes direct contact with critical residues on MA that were shown previously to be necessary for Env incorporation in virions (Figure 1H) (Tedbury and Freed 2014, Tedbury, Novikova et al. 2016).

**Figure 2.**
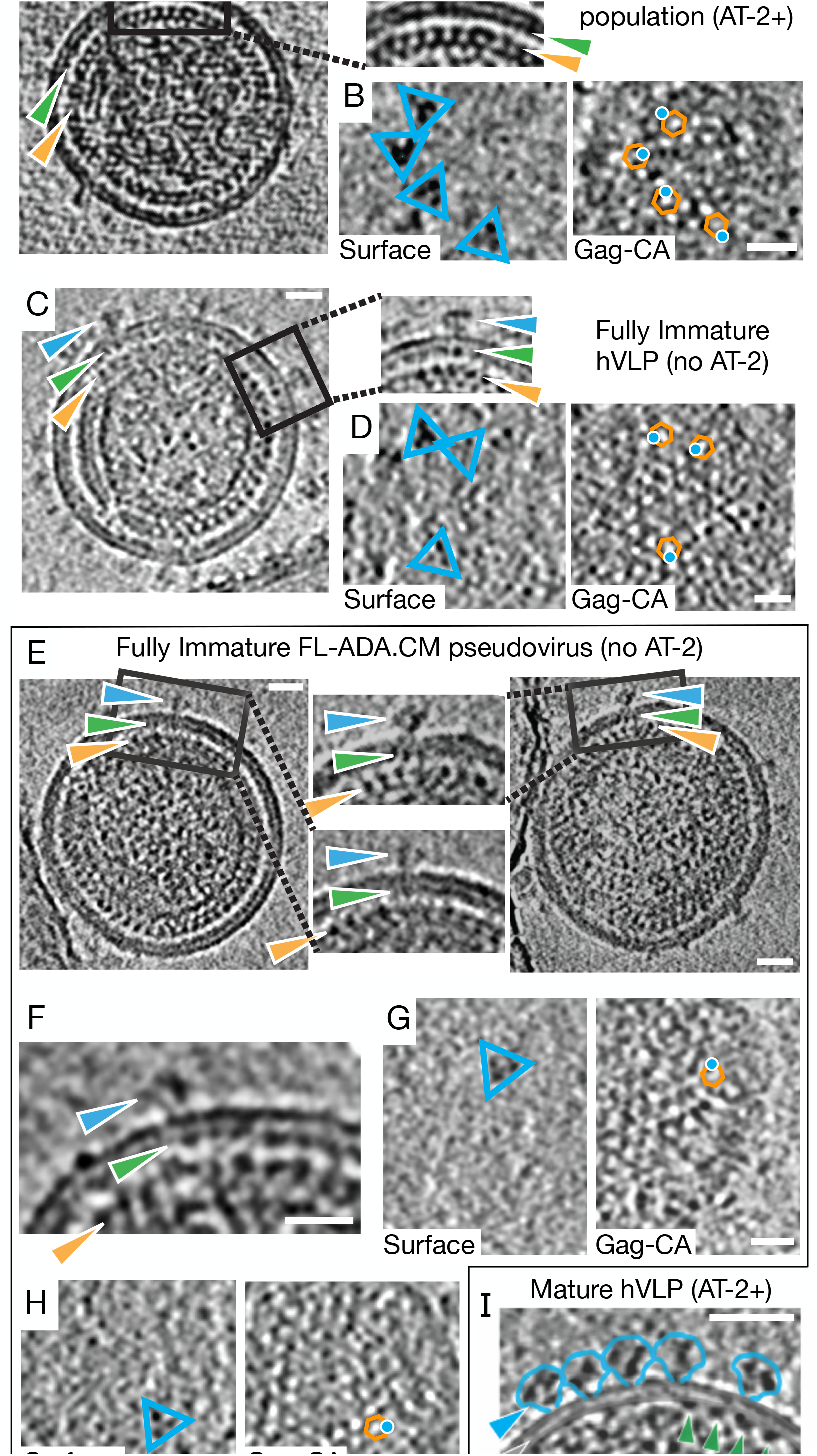
HIV Env positioning over the immature Gag lattice. **(A,B)** Representative immature sub-population of AT-2 treated ADA.CM.v4 hVLP particles. **(A)** Tomogram slices through central section showing position of Env (blue arrow) over MA (green arrow) and CA (orange arrow)**. (B)** Top view of VLP surface plane and through Gag-CA layer. Env particles are indicated by blue triangles (left panel). Right panel shows position of Env central axis (blue dot) with respect to the Gag-CA hexamer (orange hexagon) when viewing along an axis perpendicular to the plane of image. **(C,D)** Representative fully immature, non-AT-2 treated ADA.CM.v4 hVLP particles. Tomogram slices through central section **(C)**, at VLP surface plane **(D left)** and through Gag-CA layer **(D right). (E-H)** Representative fully immature, non-AT-2 treated full length ADA.CM Env bearing VLP particles. Tomogram slices through central section **(E,F)** and at VLP surface plane **(G, H left)** and through Gag-CA layer **(G,H right)**. Panels **D**, **G**, and **H** are all annotated in a similar way as panel **B**. Notably, in all of them, Env is positioned on the rim of the Gag hexamer. Black represents high density in all panels. Tomogram slice thickness is 10.32 Å in **A, B** and 8.32 Å in panels **C-H**. **(I)** Mature ADA.CM.v4 hVLP showing loss of Env (blue)/Gag-MA (green) colocalization in cross-sectional central slice. Slice thickness is 10.32 Å. Scale bars are 200Å in length.

To confirm that the observed Env positioning relative to Gag-CA in immature hVLPs is not affected by AT-2 treatment, lack of a complete C-terminal tail or incomplete processing of Gag precursor, we collected cryo-ET data of fully immature hVLPs displaying ADA.CM trimers and immature VLPs displaying full-length ADA.CM trimers, without AT-2 treatment in both cases. These fully immature VLPs were produced in the presence of protease inhibitor darunavir such that the resulting particles have greater than 95% unprocessed Gag (Figure S4). Analysis of cryo-electron tomograms from these immature VLPs having unprocessed Gag polyprotein further confirmed that the organization of the Gag polyprotein, especially the MA and CA domains, is similar to that seen in the AT-2 treated immature hVLPs (Figure 2). Additionally, Env trimers are indeed positioned along the rim of the Gag-CA lattice and not in the central hole of the Gag-CA hexamers (Figure 2D, G and H), identical to that observed in the AT-2 treated hVLP sample. Preservation of this Env-Gag colocalization in the presence of truncated as well as complete cytoplasmic tail suggests that a subset of CTD residues in the baseplate may be sufficient for mediating interactions with Gag, but the remainder of the tail may reduce incorporation into particles through steric hinderance with the Gag lattice (Buttler, Pezeshkian et al. 2018).

### Rearrangement of MA layer occurs during maturation

In immature particles, the Gag-MA layer is consistently arrayed underneath the inner membrane (Figure 1A, F and 2A, C, E, F), whereas in mature particles where the Gag polyprotein has been proteolytically cleaved, the MA layer appears fragmented (Figure 2I), indicating that MA rearranges during particle maturation, consistent with a recent report (Qu, Ke et al. 2021). Notably, Env trimers in the mature particles are no longer colocalized with the MA layer. Given this ultrastructural reorganization following maturation, it seems possible that disruption of membrane-associated Gag lattices may be a prerequisite for Env trimers to gain necessary mobility to mediate membrane fusion (Wyma, Jiang et al. 2004, Chojnacki, Staudt et al. 2012, Roy, Chan et al. 2013). These structural changes may also provide sufficient membrane pliability for remodeling during viral fusion, similar to matrix layer disruptions seen in other viruses such as influenza virus (Gui, Ebner et al. 2016).

### Sub-nanometer HIV-1 Env structure shows a conformationally variable gp41 subunit

A total of 32802 sub-volumes were used to reconstruct a C3-symmetric sub-tomogram averaged structure of hVLP-Env with local resolution ranging between 5.7 -12 Å and a global resolution of 9.1 Å (Figure 1C, 3A, B and Figure S3) (Table 1). Classification of the sub-volumes did not yield any other distinct conformational class, indicating that the hVLP-Env were in a predominantly closed, pre-fusion state. The Env ectodomain connects to the membrane via a thin tripod stalk (Figure 1C and 3B). Though the TMD can be observed spanning the membrane layer in raw tomograms (Figure 1A), no evidence of TMD is seen in the sub-tomogram averaged structure (Figure 3B), similar to the Env structure from immature hVLPs.

**Figure 3.**
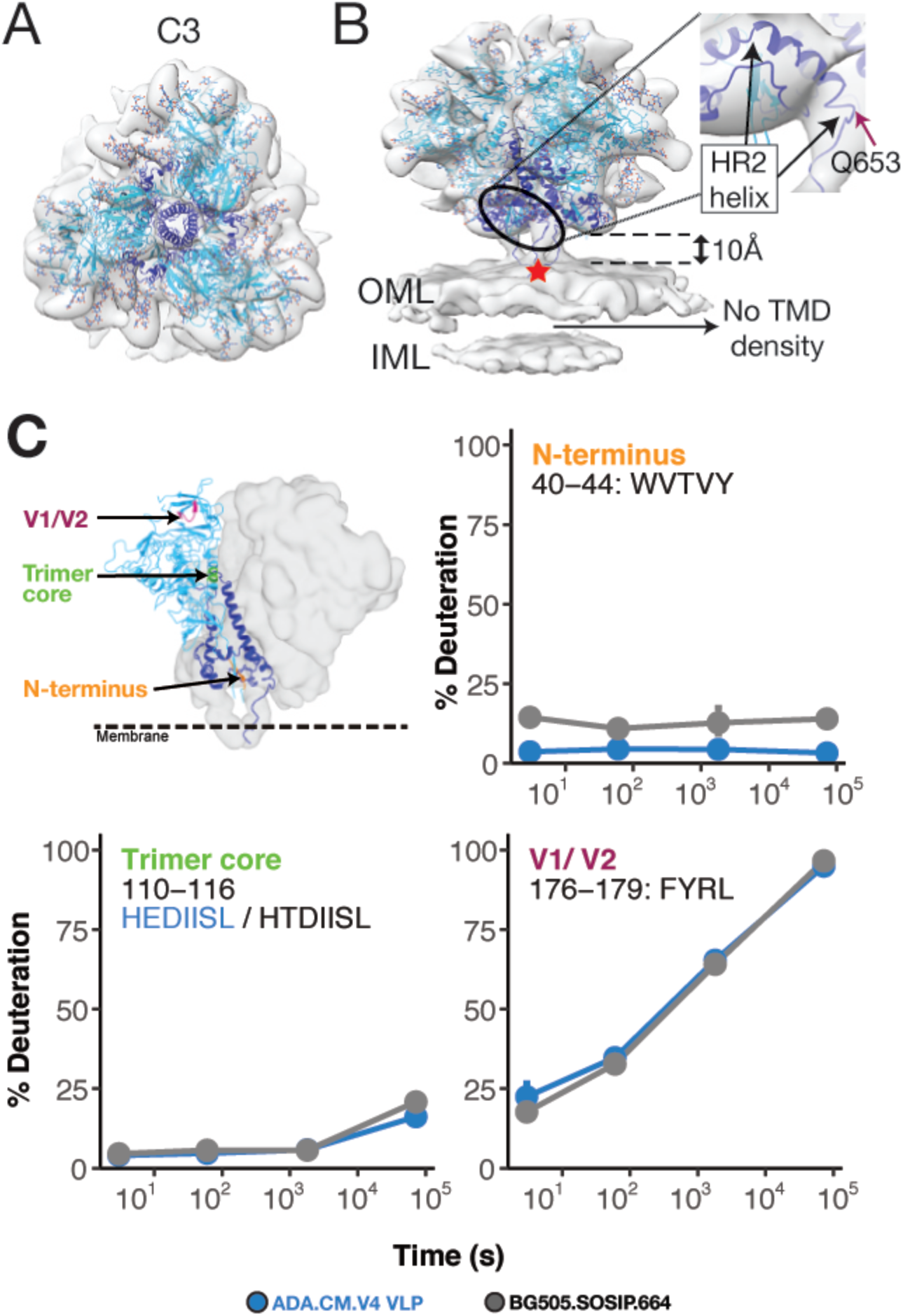
Sub-nanometer structure of hVLP-Env indicates conformational variation in HR2 helix and MPER location. Top **(A)** and side-view **(B)** of sub-tomogram averaged Env map with fitted ribbon structure adapted from full-length Env structure (PDB ID: 6ULC). Gp120 subunit is colored in cyan, gp41 in dark blue, glycans are colored as red and blue heteroatoms. Outer and inner membrane leaflets are indicated as OML and IML respectively. Zoomed inset shows the shortened HR2 helix fitted into its respective density. Red star indicates position of Asp-664 in the trimer structure (See Figure S7 for details on MPER variation). **(C)** HDX-MS of hVLP-Env shows protection of peptides that report on the closed prefusion conformation of HIV-1 Env (Guttman, Garcia et al. 2014, Guttman, Cupo et al. 2015) (see also Figures S5 and S6). Deuterium uptake for peptides reporting on trimer integrity behave similarly between hVLP-Env and BG505.SOSIP, which predominantly samples a closed prefusion conformation.

**Table 1.** Data collection and processing parameters for cryo-ET and sub-tomogram averaging.

|  |  |  |  |  |
| --- | --- | --- | --- | --- |
| Data Collection Parameters: |  |  |  |  |
| Magnification | 58000X |  |  |  |
| Voltage | 300 kV |  |  |  |
| Pixel size | 2.58 Å/pixel |  |  |  |
| Electron dose | 64-68 e<sup>-</sup>/Å<sup>2</sup> |  |  |  |
| Defocus Range (µm) | -2.5 to -5.0 |  |  |  |
| Total number of tilt-series | 526 |  |  |  |
| Final number of tilt-series | 423 |  |  |  |
| Data Processing Parameters: |  |  |  |  |
|  | hVLP-Env<br>C3 symmetry<br>(sym) | hVLP-Env<br>C1 sym | Immature<br>hVLP-Env<br>C3 sym | Immature<br>hVLP-Env<br>C1 sym |
| Initial number of sub-volumes | 63592 | 63592 | 1520 | 1520 |
| Final number of sub-volumes | 32802 | 29074 | 1051 | 1059 |
| Map resolution at 0.143 FSC | 9.13 Å | 10.67 Å | 31.35 Å | 34.3 Å |

Env copy number on individual mature hVLP particles ranged from 8-82 Env and the trimers appeared to be randomly distributed with no apparent clustering or predisposition to forming close interactions with other trimers (Figure 1B). The hVLPs have a predominantly spherical morphology in our sample with an average diameter of ∼100nm. Thus, by calculating the surface area of the VLPs as a sphere and the Env trimer as an equilateral triangle with sides of 13nm length, we can determine that an average hVLP can accommodate up to 429 Env trimers (31400nm^2^/73.18nm^2^) without clashing. Thus, although hVLPs display high levels of Env compared to wild-type HIV, the number of surface Env trimers present on hVLPs was much lower than the theoretical maximum, so Env trimers were not cramped and have sufficient surface area for mobility. Furthermore, the sub-volumes used in the final reconstruction of hVLP-Env were also selected to exclude neighboring Env in close proximity to avoid any influence due to unlikely clashing.

To gain complementary structural insight into membrane embedded Env, we conducted hydrogen-deuterium exchange mass-spectroscopy (HDX-MS) analysis of Env from intact hVLPs. This approach samples the entire population of Env in our specimen. HDX-MS analysis showed that key fiducial peptides which report on trimer integrity (Guttman and Lee 2013, Guttman, Garcia et al. 2014), were well-protected in hVLP-Env (Figure 3C), confirming its closed, pre-fusion conformation. Moreover, the HDX-MS spectra were unimodal, consistent with a homogenous conformational population of Env. Relative to standard HDX-MS experimental approaches, we were able to improve peptide coverage by performing deuterium exchanges under native conditions but solubilizing the surface protein using detergent under quench conditions (Figure S5). Comparison of hVLP-Env with BG505.SOSIP showed that cognate peptides throughout the trimers had similar exchange profiles (Figure S6), indicating that they both have comparable global conformations. However, the hVLP-Env trimer on virions exhibited greater exchange protection, indicating it is in general more structurally ordered than the engineered soluble trimer (Figure 3C and S6). Indeed, in terms of gp120 subunit organization, hVLP-Env has a global architecture that closely resembles previously published Env structures (Kwon, Pancera et al. 2015, Ward and Wilson 2015, Lee, Ozorowski et al. 2016, Pan, Peng et al. 2020). This is confirmed by good agreement seen in fitting the atomic structure of detergent extracted full-length Env (PDB ID: 6ULC) into the sub-tomogram averaged map (Figure 3A, B).

### Differences in gp41’s terminal HR2 helix and stalk influence MPER surface exposure

Significant differences relative to available structures are evident, however, in the gp41 subunit of hVLP-Env. In nearly all known Env structures, the HR2 helix has a long, rod-like conformation with Asp664 forming its distal tip (Kwon, Pancera et al. 2015, Ward and Wilson 2015). In contrast, in hVLP-Env, Gln-653 forms the distal tip of the HR2 helix, making the helix length 10 amino acids shorter (Figure 3B). The remaining residues in the C-terminal portion of HR2 helix bend and form a thin stalk connecting the ectodomain to the membrane (Figure 3B). Apart from hVLP-Env, this bend in HR2 helix has only been observed in the full-length, detergent-solubilized Env structure derived from strain 92UG037.8 (Pan, Peng et al. 2020). In the presence of membrane, we observe that the thin stalks, corresponding to Env residues 654-664, form a tripod that elevates the ectodomain ∼10 Å above the membrane (Figure 3B). As a result, the bulk of the MPER (Membrane-Proximal External Region) (residues 660-683), which is a desirable vaccine target of some of the most broadly neutralizing HIV-1 antibodies isolated to date (Gray, Madiga et al. 2009, Rantalainen, Berndsen et al. 2020), is primarily embedded within the membrane in our structure.

The complete MPER peptide is unresolved in all reported structures of trimeric Env, owing to its membrane proximal location and/or possible variations in its conformation. Current knowledge of MPER structure is only derived from constructs without the Env ectodomain included (Fu, Shaik et al. 2018, Piai, Fu et al. 2020). In a recently published structure of Env from BaL-1 virions, the MPER was indicated to form part of the stalk connecting Env ectodomain to the membrane (Li, Li et al. 2020), which contrasts with the membrane embedded MPER location identified in our structure (Figure S7). However, the embedded nature of the MPER in our map is in agreement with that inferred from other solubilized full-length Env structures and biophysical studies on binding of MPER-targeted antibodies to HIV-1 Env (Kim, Sun et al. 2011, Lee, Ozorowski et al. 2016, Wang, Kaur et al. 2019, Rantalainen, Berndsen et al. 2020). Analyzing Env structures from different HIV strains shows that the general position of the bulk of the ectodomain is ∼10-12 Å above the interpreted membrane surface in all cases (Figure S7). However, differences arise in the position of HR2 helix and subsequent MPER sequence leading to variability in MPER presentation and accessibility above the membrane amongst HIV-1 strains.

### A flexible gp41 stalk allows Env orientational freedom and MPER epitope exposure

In our tomographic data, we observe tilting of hVLP-Env with respect to the membrane (Figure 4A), which leads to a lower density level for the stalk in our sub-tomogram averaged map (Figure 4B). Thus, the tripod stalk of Env is flexible in its membrane embedded state, allowing differential sampling of MPER and other membrane-proximal epitopes. Although tilting or lifting of Env from the membrane has been suggested based on bound complexes with MPER targeted antibody 10E8 (Lee, Ozorowski et al. 2016, Rantalainen, Berndsen et al. 2020) and gp120-gp41 interface targeting antibody 35O22 (Huang, Kang et al. 2014), our results provide the first direct evidence of Env tilting on native membrane in an unliganded state.

**Figure 4.**
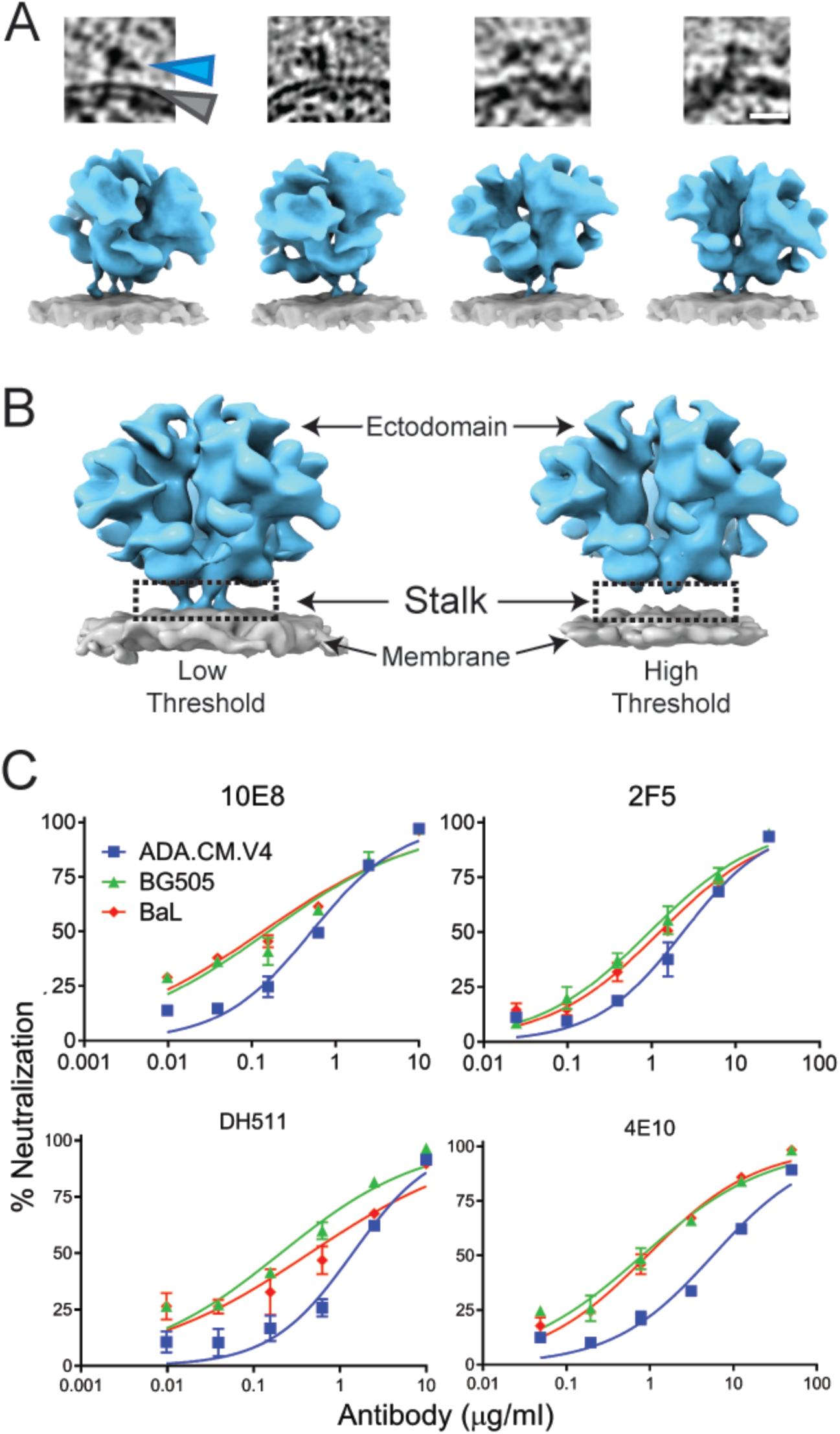
Flexible gp41 stalk in HIV Env leads to variation in MPER epitope accessibility. **(A)** Top: Tomogram slices showing examples of tilted Env on membrane surface. Scale bar equals 100 Å. Bottom: Surface rendering of modeled Env structure (blue) in tilted orientations on membrane (grey) as seen in the tomograms. (**B)** Env density map rendered at low and high thresholds, respectively, showing loss of density in the stalk region (black rectangle). **(C)** Comparison of neutralization effect on different HIV-1 strains by MPER-directed bnAbs. Also see Figure S8.

Sensitivity to MPER broadly neutralizing antibodies (bnAbs) has been shown to be impacted by Env stability in its prefusion state (Kim, Leaman et al. 2014), with increased neutralization observed after cellular receptor engagement (Frey, Peng et al. 2008, Chakrabarti, Walker et al. 2011). Based on the structural data discussed above, we envision that local structure of HR2, height of the Env ectodomain above membrane, and propensity of Env tilting have additional impact on Env sensitivity or resistance to MPER bnAbs. To test this hypothesis, we conducted comparative neutralization assays using MPER bnAbs against HIV-1 strains whose Env structures have been determined. As anticipated from the structural data, amongst the three viruses, ADA.CM hVLP, BaL-1 and BG505, ADA.CM was overall more resistant to MPER bnAbs (Figure 4C and Table S2). Neutralization and mutational analysis of proximal residues in ADA.CM that contribute to trimer stability (Figure S8) showed modest individual effects but none were individually responsible for ADA.CM’s enhanced resistance to MPER bnAbs (Figure S8 and Table S3).

### Structural variation in gp41’s HR1-C helical bundle suggests a conformationally variable central core

Another prominent difference in the gp41 subunit of hVLP-Env is absence of complete density in the averaged structure for the central C-terminal HR1 helices (HR1-C) (Figure 5A). The HR1-C helices comprised of amino acid residues 570-595 have been a hallmark of high resolution Env SOSIP structures with the central trimeric bundle composed of a HR1-C helix from each of the gp41 domains. However, in our structure, only partial density for the HR1-C helix can be observed corresponding to gp41 residues 570-577 and 582-595 (Figure 5A). Classification of Env sub-volumes with a tight cylindrical mask for the gp41 region did not yield any subset with higher density for HR1-C helices in our data. Considering that our sub-tomogram averaged map has a local resolution range for the ectodomain between 5.7-12 Å (Figure S3), it is clear from comparisons with other cryo-EM maps at similar resolutions, that the absence of complete HR1-C helix density in our structure is not an effect of map resolution (Figure S9).

**Figure 5.**
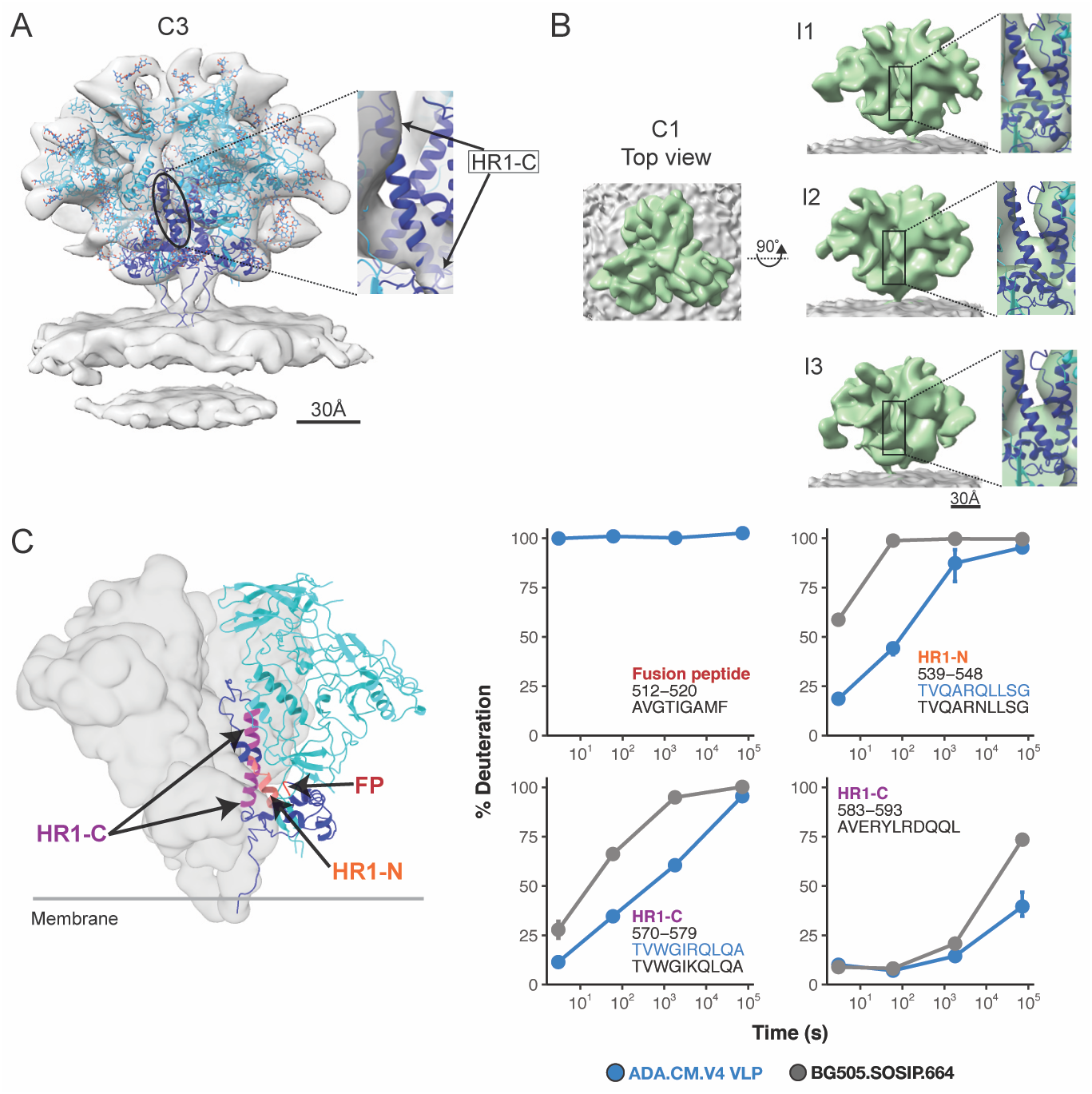
Conformational variation in central helical bundle (HR1-C) of hVLP-Env. **A)** Side-view of sub-tomogram averaged Env map with fitted ribbon structure and colored similar to Figure 1. Zoomed inset shows fit of HR1-C helix into its respective density (See also Figure S9). **B)** Top and side views of an asymmetric reconstruction of hVLP-Env with the three interfaces (I1, I2 and I3) of Env trimer shown along with magnified insets of fitted HR1-C helix. Black rectangle indicate the central core of the trimer where the HR1-C helix lies. See Figure S3 for resolution-related details. **C**) HDX-MS deuterium uptake plots for fusion peptide (FP) and HR1 peptides (see also Figures S5 and S6). Surface rendering of Env was calculated using the fitted atomic model of hVLP-Env in panel A without glycans. Full FP region is not observed in hVLP-Env structure due to flexibility.

To ascertain whether the observed HR1-C density was affected by applied symmetry to the map, we calculated an asymmetric structure of hVLP-Env at 10.7 Å resolution (Figure S3 and Table 1). The asymmetric structure (Figure 5B) appears overall similar to the three-fold symmetrized map (Figure 5A). Partial density for the HR1-C helices is consistently present across all three protomers in the asymmetric structure, with near-complete density for HR1-C observed in one protomer (Figure 5B).

HDX-MS analysis provides additional insight into the state of this key structural motif. Peptides in HR1-C helix (residues 570-579 and 583-593) are more protected in hVLPs than in BG505.SOSIP (Figure 5C), indicating increased backbone hydrogen bonding interactions in corresponding residues. Taken together, the sub-tomogram averaged structures and HDX-MS analysis shows that the peptides in HR1-C region exist in a stable, ordered conformation in hVLP-Env even though they do not rigidly conform to Env’s global three-fold structural symmetry.

### An extensive and heterogenous glycan shield in hVLP-Env protects critical epitopes

The hVLP-Env structure displays distinguishable density features for N-linked glycans that decorate the ectodomain surface. In contrast to most available structures where density near to glycosylation sites often covers only the core GlcNAc sugars, in our unliganded hVLP-Env structure, extended density for glycan moieties is observed across the protein. Resolved density for multiple canonical glycan positions in gp120 and gp41 subunits extends beyond the range of current structural models, revealing the extent of the ‘glycan shield’ (Figure 6A) (Wei, Decker et al. 2003). Glycans surrounding several key bnAb epitopes can be identified in the unliganded Env map, which reveal their orientations in the natural, antibody-naïve state so as to occlude underlying epitopes.

**Figure 6.**
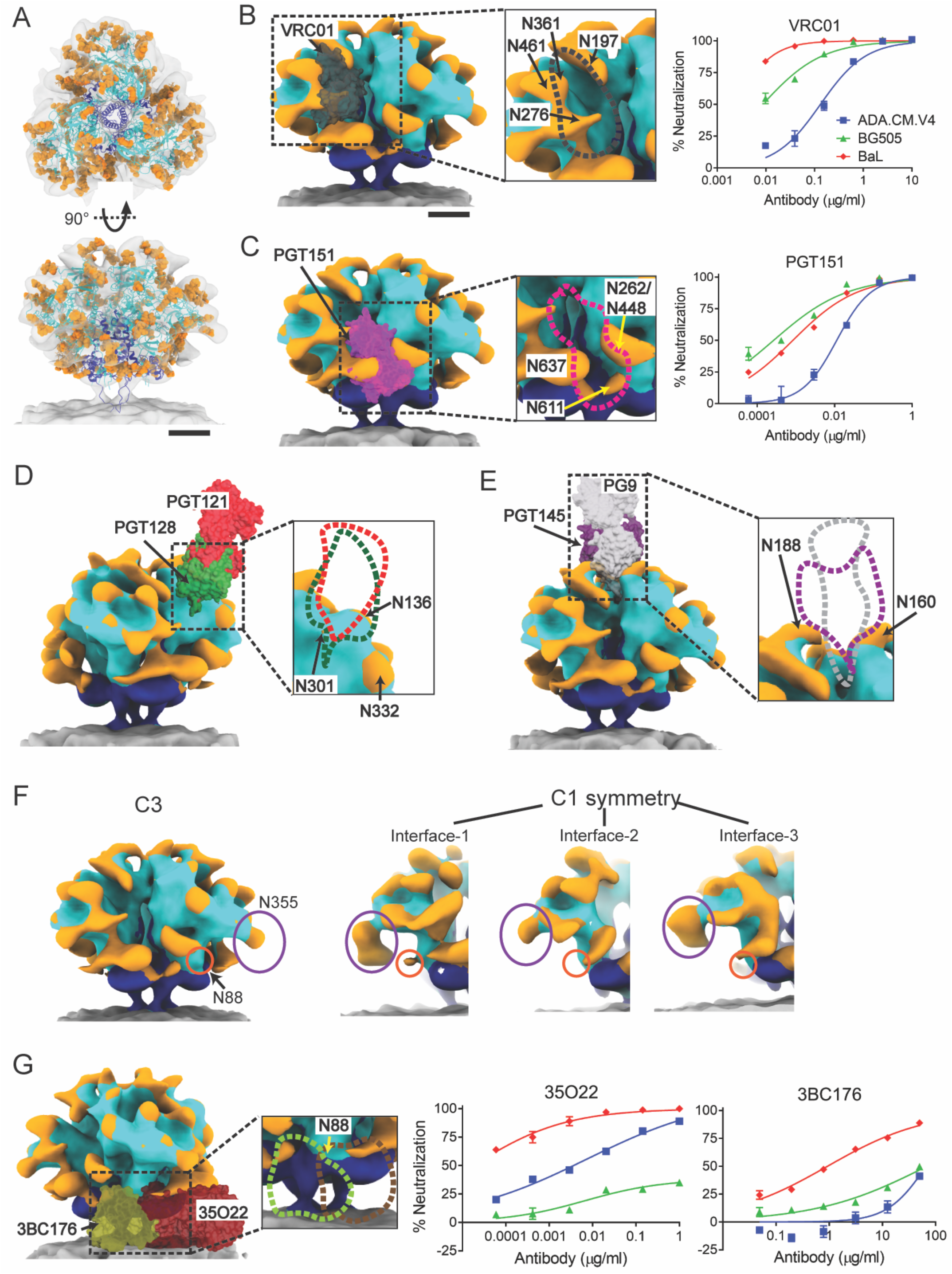
Presence of an extensive glycan shield in hVLP-Env. **(A)** Top and side view of hVLP-Env with glycans rendered as orange ball-stick structures. Gp41 colored in dark blue and gp120 in cyan. **(B-E and G)** Binding positions of eight bnAbs – VRC01 (dark grey), PGT151 (pink), PGT128 (green), PGT121 (red), PG9 (light grey), PGT145 (purple), 35O22 (brown) and 3BC176 (green) mapped onto hVLP-Env surface. Gp120, gp41, glycan surfaces and membrane are colored in cyan, dark blue, orange and grey respectively. Zoomed-in sub-panel outlines interaction surfaces corresponding to each bnAb. Glycans in the vicinity of bnAb binding sites are labeled. Neutralization of HIV-1 strains, ADA.CM.v4, BG505 and BaL in presence of bnAb are shown in B, C and G. **F)** Differential glycosylation among trimer subunits in hVLP-Env represented using glycans at N88 (orange circle) and N355 (purple oval). Surface rendering of C3-symmetrized map on left followed by the three protomer interfaces from the C1 symmetry hVLP-Env map. Scale bars equal 30 Å.

At the CD4 receptor binding site (CD4bs) in hVLP-Env, overlaying the previously determined Fab (Fragment antigen binding) structure of VRC01 (CD4bs bnAb) (Stewart-Jones, Soto et al. 2016) in its binding orientation, highlights possible clashes with nearby glycans N197, N276, N361 and N461 (Figure 6B). These glycans have also been shown to overlap with the VRC01 epitope using molecular dynamic studies, but previously determined structures using SOSIPs resolved only short glycan chains at these sites (except for N276) resulting in minimal apparent glycan interactions with the bnAb (Stewart-Jones, Soto et al. 2016). In hVLP-Env, extended density for these multiple glycans can be seen to clearly deter binding of CD4 epitope-directed nAbs (Figure 6B). These observations are corroborated in VRC01 based neutralization assays which show that hVLP-Env is substantially more resistant than BaL-1 Env (Figure 6B and Table S2), whose structure lacks extended glycosylation around this site (Li, Li et al. 2020). Indeed, removal of N276 and N461 glycans has been demonstrated to facilitate engagement of germline precursor antibodies for VRC01-like nAbs that normally do not interact strongly with natively glycosylated Env (McGuire, Hoot et al. 2013).

Similarly, long-range extended densities for N-linked glycans are observed in the gp120 domain corresponding to residues N262 and N448 as well as in the gp41 domain corresponding to residues N611 and N637. The close spacing of N262/N448 to N611 likely orders these glycans to be visible in the averaged map (Figure 6A, C). Collectively, these glycans form a protective barrier over the fusion peptide (FP) in hVLP-Env (Figure 6C). Env’s FP is essential for viral cell entry but is surprisingly exposed to solution and highly dynamic by HDX-MS analysis (Figure 5C). Mapping the Fab structure of FP targeting bnAb, PGT151 (Lee, Ozorowski et al. 2016), onto its corresponding epitope on hVLP-Env shows how the glycans would need to be displaced in order for the antibody to bind (Figure 6C). Though hVLPs are neutralized by PGT151, they are relatively resistant compared to other strains (Figure 6C and Table S2), suggesting that glycan occlusion may impact neutralization by FP targeted bnAbs as has been reported (Stewart-Jones, Soto et al. 2016).

Notably, we do not observe strong clashes with glycans at the V3-loop and V1/V2 loop antibody binding super-sites. Overlaying the Fab structures of V3-loop targeting antibodies PGT121 (Garces, Lee et al. 2015) and PGT128 (Lee, de Val et al. 2015), shows that the Fabs fit snugly between glycans in that region, particularly N332, N136 and N301 (Figure 6D). Presence of an extended N301 glycan might provide some hindrance to binding of PGT128 Fab, but we only observe minimal ordered density for glycosylation at N301 in our map, which limits further analysis. Similarly, when the Fab structures of V1/V2-loop targeting antibodies PG9 (Wang, Gristick et al. 2017) and PGT145 (Lee, Andrabi et al. 2017) were mapped onto the sub-tomogram averaged Env map, no clashes were observed with nearby glycans at positions N160 and N188 (Figure 6E). Antibodies to the V3-loop and V1/V2 loops, generally rely on extensive contacts with neighboring glycans for their activity. As can be seen from our analysis, absence of occlusion of their respective binding sites by the surrounding glycans might be an additional factor for these sites to be targeted efficiently by antibodies (Andrabi, Voss et al. 2015, Moyo, Kitchin et al. 2020).

Extensive N-linked glycosylation densities also persist in the asymmetric structure of hVLP-Env. Strikingly, we observe variations in glycan densities between different protomers, indicating heterogeneity in glycan occupancy. For example, the N355 glycan is present at all three protomers in the asymmetric Env reconstruction but its density varies with respect to size and orientation at this site (Figure 6F). Similarly, the conserved N88 glycan, which forms part of the epitope of bnAb 35O22 (Huang, Kang et al. 2014) and borders that of 10E8 (Lee, Ozorowski et al. 2016), can only be observed in one protomer but not in others (Figure 6F) and is absent in the C3-symmetrized map. In mass spectrometry analysis of hVLP-Env, the glycan site at N88 along with several others, appear to be occupied by a heterogenous population of complex sugars (Table S4). Although the observed heterogeneity in glycan processing of hVLP-Env (Table S4) is similar to previous reports (Guttman, Garcia et al. 2014, Cao, Pauthner et al. 2018, Berndsen, Chakraborty et al. 2020), our comparative structural analysis on asymmetric and symmetric hVLP-Env directly resolves glycan heterogeneity within protomers of the same Env trimer.

When comparing HIV bnAbs affected by both the presence of glycan and membrane proximity, binding of bnAb 35O22 (Huang, Kang et al. 2014) is highly dependent on the presence of N88 glycan whereas 3BC176 binding (Lee, Leaman et al. 2015) is partially inhibited by it. Mapping binding orientation of both these Fabs onto hVLP-Env results in steric clashing of the Fab regions with the membrane surface (Figure 6G). Nevertheless, hVLPs are effectively neutralized by 35O22 but show relative resistance to 3BC176 (Figure 6G and Table S2), presumably due at least in part to 3BC176’s lower binding capacity (Lee, Leaman et al. 2015). Thus, a combination of the heterogenous presence of N88 glycan and tilting of hVLP-Env on the membrane is consistent with the differential access of 35O22 and 3BC176 to their epitopes on membrane embedded Env.

### Conclusions

In this study, we have analyzed the structure of HIV particles displaying functional, membrane-bound Env. The ADA.CM Env trimers used are highly stable in a membrane environment, although they notably do not form well-ordered gp140 trimers in solution using the typically employed “SOSIP” mutations (Leaman and Zwick 2013). This is true for the majority of Env sequences, that require extensive stabilizing modifications in order to encourage formation of stable ectodomain trimers (Rawi, Rutten et al. 2020). Thus, the structural details discussed here may reflect features of Env strains that are not compatible with SOSIP modifications and thus have been resistant to structural characterization so far.

In hVLP-Env, we find substantial evidence that key parts of gp41 including the HR1 central helices and flexible stalk are not rigidly fixed relative to the rest of the trimer. Indeed, R.M.S.D. calculations among static high-resolution structures of full-length Env and SOSIP structures show higher deviation in the gp41 domain compared to gp120 domain (Table S5), despite the strains having greater than 75% sequence similarity (Table S6). Likewise, DEER spectroscopy studies of SOSIP trimers have also shown a higher degree of variability in the gp41 domain (Stadtmueller, Bridges et al. 2018), further substantiating the flexible nature of gp41 as inferred from our structures.

Our data also demonstrate that even in membrane-associated Env, the fusion peptide (FP) is highly dynamic and exposed to solvent. Exposure of such a functionally critical and conserved component seems counter-intuitive, but the dense clustering of glycans around the FP proximal region likely confers a degree of steric protection. However, breaches in the glycan shield, revealed by heterogeneity in glycan density in this region in our structures, can still offer access to the FP by bnAbs (Kong, Xu et al. 2016, Lee, Ozorowski et al. 2016). Moreover, in cases where glycans form part of bnAb epitopes, heterogeneity in glycosylation provide mechanisms exploited by HIV-1 as a means of increasing epitope variability and facilitating evasion of antibody responses. Finally, the extent of ordered density between adjacent glycans in hVLP-Env suggests that the glycans are interacting in a stable manner, consistent with recent reports (Stewart-Jones, Soto et al. 2016, Berndsen, Chakraborty et al. 2020).

By imaging Env in a membrane-bound context on VLPs, we found that, like other type I fusion proteins such as SARS-CoV-2 S trimers (Ke, Oton et al. 2020), HIV-1 Env ectodomain sits atop a flexible stalk that affords it considerable freedom of motion to tilt relative to the membrane. This likely plays an important mechanistic role by affording fusion proteins the flexibility to bind receptors and refold during membrane fusion. The flexible stalk also impacts the accessibility of key epitopes that are sterically hindered on the membrane-facing side of the trimer ectodomain such as the FP, gp120/gp41 interface, and membrane-associated MPER. Thus, our structural analyses revealed previously uncharacterized Env features that impact presentation of prime epitopes that can help explain differences in neutralization sensitivity across diverse HIV-1 strains. Lastly, our structural analyses of Env in immature viral particles, shows that Env is situated directly over two-fold symmetric contacts of the Gag lattice, providing the first structural evidence of a direct physical Env-Gag interaction, helping to inform models for virion particle assembly and Env incorporation. Taken together, our results advance understanding of HIV-1 Env in the context of virion assembly, maturation and conformational sampling, revealing insights that are generally unavailable through structural studies on recombinant proteins alone.

## Limitations

The ADA.CM.v4 hVLPs, which is the primary sample used in our experiments, display Env that has a C-terminal truncation of 102 amino acids. This leaves about 40 amino acids of the CTD on the Env protein, sufficient to form part of the baseplate revealed in a recent NMR structure (Piai, Fu et al. 2020). The loss of the rest of the tail could potentially have implications in Env recruitment and assembly in the VLPs. The hVLP sample was also treated with aldrithiol-2 (AT-2), a mild oxidizing agent, which inactivates HIV by causing the zinc finger domain of the nucleocapsid subunit to release zinc (McDonnell, De Guzman et al. 1997). However, it does not act directly on the Env protein, which our data and others have showed is intact in terms of structure, antigenicity, and cell receptor-mediated membrane fusion activity (Rossio, Esser et al. 1998, Rutebemberwa, Bess et al. 2007).

Our analyses of non-AT2 treated VLPs, produced in the presence of the HIV protease inhibitor darunavir and displaying truncated as well as full-length Env, showed that AT-2 treatment did not affect direct Env interactions with the underlying Gag lattice in the immature VLPs. In the mature hVLPs, the processed Gag core is not in direct contact with surface Env; however, we cannot rule out an indirect effect of AT-2 on nucleocapsid and the core. Lastly, a minor population of unprocessed Env (gp160 trimers) may exist, particularly on the VLPs bearing full-length Env, which cannot be distinguished from mature Env (gp120-gp41 heterotrimers) at the current resolution of our data. Despite the caveats above, in the range of immature particle examples we have examined, Env-MA-Gag interactions remain notably consistent, thus underscoring a persistent coupling between these key structural components of HIV.

## Acknowledgements

We thank the University of Washington Arnold and Mabel Beckman Cryo-EM Center and the School of Pharmacy Mass Spectrometry Center for data collection time and support. This work was supported by National Institutes of Health grants S10OD032290 and NIH NIGMS and NIAID grants R01GM099989 and R01AI140868 to K.K.L., T32 GM008268 to M.A.B., and R01AI143563 to M.B.Z., the James B. Pendleton Charitable Trust (M.B.Z.), and by the University of Washington’s Proteomics Resource (UWPR95794). A grant from the Bill & Melinda Gates Foundation (OPP1126258) also supported portions of this work.

## Author Contributions

V.M.P.: Conceptualization, Formal analysis, Investigation, Validation and Writing. D.P.L.: Investigation, Formal Analysis. K.N.L.: Investigation, Formal Analysis. J.T.C: Investigation, Formal Analysis. M.A.B.: Investigation. E.A.H.: Investigation. M.B.Z.: Resources, Writing review, Funding acquisition. K.K.L.: Conceptualization, Writing, Funding acquisition.

## Data Availability

Density maps with corresponding atomic models have been deposited with accession codes EMD-XXXX, EMD-XXX, EMD-XXX, EMD-XXXX and PDB IDs XXXX and XXX.

## Experimental Methods

### hVLP expression and purification

Stable cell line expressing high levels of HIV Env, ADA.CM.v4, was described previously (Stano, Leaman et al. 2017). A stable cell line expressing Env BG505 was generated using a similar method, namely by transduction of HEK293T cells using a lentiviral vector (pLentiIII.BG505.755*) containing the *env* gene BG505 with a stop codon at position 756. Transduced cells were subjected to two rounds of sorting by FACS using fluorescently labeled quaternary neutralizing antibody PGT145 and non-neutralizing CD4BS antibody b6, then gating cells for PGT145^high^ b6^low^. Env copy number on BG505 VLPs was notably lower than on ADA.CM.v4 hVLPs. Additionally, BG505 VLPs were also more fragile, which precluded extensive cryo-ET data collection and HDX-MS analysis.

To generate hVLPs, cells were transfected using Env-deficient HIV-1 backbone plasmid pSG3ΔEnv (NIH ARRRP) and 25 kDa PEI (Polysciences) as a transfection reagent. Immature hVLPs and full-length ADA pseudovirus were produced by adding the protease inhibitor darunavir (NIH AIDS Reagent Program) to transfected cells at the same time as the DNA and PEI to a final concentration of 2 µM. Supernatants were harvested 3 days after transfection and cleared by centrifugation at 3,000 × *g* for 15 min. hVLPs were pelleted at 80,000 × *g* for 1 h and resuspended 100-fold concentrated in PBS. VLPs were separated from cellular debris using iodixanol density gradient centrifugation. Here, concentrated hVLPs were overlaid on a 9.6 to 20.4% iodixanol (Optiprep; Sigma) gradient, formed by layering iodixanol in 1.2% increments, and centrifuged at 200,000 × *g* for 1.5 h at 4°C in an SW41Ti rotor (Beckman). Fractions (1 ml each) were collected starting from the top and fractions 6-9 were saved and pooled. Purified hVLPs were brought up to 15 ml with PBS and concentrated to ∼0.2 ml using a 100 kDa MWCO Amicon centrifugal filter (Millipore). VLPs were inactivated by adding aldrithiol-2 (AT-2) to a final concentration of 2.5 mM and samples were incubated at RT for 2 h. The volume was again increased to 15 ml with PBS and hVLPs were concentrated using a 100 kDa MWCO centrifugal filter to 250-fold the concentration in the original transfection supernatant.

### Cryo-ET sample preparation and tilt-series acquisition

Purified AT-2 treated hVLP samples were mixed with 10nm gold beads (Aurion BSA Gold Tracer 10nm) at a ratio of 15:1 (v/v). Using a Vitrobot Mark IV (FEI Co.), 3 μl of this mixture was applied to glow discharged, C-Flat 1.2/1.3, 200mesh or 400mesh grids (Electron Microscopy Sciences), blotted for 4-5 seconds and plunge frozen in liquid ethane. For the non-AT-2 treated immature ADA.CM VLPs and immature full-length ADA VLPs, purified sample was mixed with 10nm gold beads at a ratio of 10:1; 3 ul of this mixture was applied to glow-discharged 400 mesh Cu Quantifoil R 1.2/1.3 grids (Electron microscopy sciences) and plunge frozen in liquid ethane. Vitrobot was maintained at 4°C and 100% humidity during all these experiments.

For the AT-2 treated hVLPs, frozen grids were imaged using a 300 kV Titan Krios with a Gatan K2 direct electron detector and GIF energy filter with slit width of 20 eV. Tilt-series were collected in a dose-symmetric tilting scheme from −54° to +54° or from −48° to +48° with a step size of 3° using Leginon (Carragher, Kisseberth et al. 2000) or SerialEM softwares (Mastronarde 2005). Tilt-series were collected in counting mode at a magnification of 53000X, corresponding to a pixel size of 2.58 Å per pixel. The total dose per tilt series was ∼64-68 e^-^/Å^2^. A total of 526 tilt-series were collected across multiple sessions.

For the non-AT-2 treated immature ADA.CM.v4 hVLPs and immature full-length ADA.CM pseudovirus, frozen grids were imaged on a 300 kV Titan Krios equipped with a Gatan K3 direct electron detector and a GIF energy filter with a slit width of 20 eV. Tilt series were collected from −60° to +60° with a step size of 3° using SerialEM (Mastronarde 2005). A dose-symmetric tilt scheme was applied between −48° and +48°, and the remaining angles were collected with a bidirectional approach. Images were collected in super-resolution mode at a nominal magnification of 64,000X, corresponding to a pixel size of 0.6932 Å/pixel. The total cumulative dose was 94 e-/Å2. For the non-AT-2 treated immature full-length ADA.CM VLPs, a total of 39 usable tilt-series were collected. For the non-AT-2 treated immature ADA.CM.v4 hVLPs, 52 usable tilt-series were collected.

For the non-AT-2 treated immature full-length ADA.CM pseudovirus, additional data was also collected on a 200kV Glacios microscope with a K2-Summit direct electron detector. A total of 17 tilt-series were collected with a nominal magnification of 28000X, corresponding to a pixel size of 1.4975 Å/pixel. Tilt series were collected from −60° to +60° with a step size of 3° using SerialEM with upto 45° in dose-symmetric tilt scheme (Mastronarde 2005). A total dose of 110 e-/Å^2^ was applied to each tilt-series.

### Tomogram reconstruction

For the AT-2 treated hVLP dataset, tilt-series image frames were corrected for beam-induced motion using motioncor2 (Zheng, Palovcak et al. 2017). Using batch tomography via the Etomo interface in IMOD software package (Kremer, Mastronarde et al. 1996, Mastronarde and Held 2017), tilt-series were processed for tomogram generation using standard procedures. Tilt series images were aligned using gold bead markers and aligned tilt-series were used to generate a three-dimensional volume using weighted back-projection method. The final tomograms were rotated, binned, and low pass–filtered for visualization. Tilt-series with non-optimal alignments were discarded. In the end, 423 tilt-series were selected for further processing.

For immature ADA.CM VLPs and immature full-length ADA VLPs, Images were motion-corrected using motioncor2 (Zheng, Palovcak et al. 2017) and tomograms were reconstructed and CTF corrected in EMAN2 using default parameters (Chen, Bell et al. 2019).

### Sub-tomogram averaging of hVLP-Env

#### Initial model generation

Eight tilt-series were randomly selected. CTF estimation and correction for these were done using the Ctfplotter program (Xiong, Morphew et al. 2009). Tomograms were then generated for these CTF-corrected tilt-series in IMOD at the unbinned pixel size (Kremer, Mastronarde et al. 1996, Mastronarde and Held 2017). The aligned tilt-series, reconstructed tomograms, file listing the tilt angles and electron dose values were imported into Relion software suite (Scheres 2012) according to its conventions (Bharat and Scheres 2016). Using the 3dmod graphical user interface (Kremer, Mastronarde et al. 1996), 2067 Env particles on the surface of hVLPs were manually picked. The picked particle coordinates were imported into Relion (Scheres 2012) and corresponding sub-volumes extracted. These were then subjected to 3D classification with a spherical mask covering small parts of the membrane using C1 and C3 symmetry with no initial model provided. Both the C1 and C3 symmetry classifications resolved a 3D class that looked similar to the expected structure of HIV-1 Env. The 3D class from the C3 symmetry run showing clear Env density contained a total of 811 sub-volumes and was selected as initial model.

#### For C3-symmetrized hVLP-Env structure

Tilt-series were imported into EMAN2’s sub-tomogram averaging pipeline (Chen, Bell et al. 2019). 1k X 1k tomograms were generated within EMAN2 using default parameters. The binned tomograms were then used for semi-automated particle picking in PEET software (Nicastro, Schwartz et al. 2006). Consolidated particles from the semi-automated picking were further curated manually using the 3dmod interface (Kremer, Mastronarde et al. 1996) to remove wrongly positioned particle points such as those that were in the membrane or on the inside of VLPs. Curated particle coordinates (63592 particles) were then imported into EMAN2 through the e2spt_boxer.py interface (Chen, Bell et al. 2019). CTF-estimation for all the tilt-series and subsequent CTF-corrections were carried out within EMAN2. Sub-volumes were initially extracted at 4xbinning corresponding to a pixel size of 10.32 Å per pixel. Sub-tomogram refinement was carried using a spherical mask including Env on surface and a part of the membrane. The 3D structure previously generated in Relion (Scheres 2012, Bharat and Scheres 2016) was used as initial model after low pass filtering to 60 Å. Sub-volumes were then re-extracted at 2X binning using the particle orientations from the 4X binned refinement output. Sub-tomogram refinement was repeated using the 2X binned data with a cylindrical mask covering only the ectodomain and outer membrane layer. Particle orientations were locally refined within a 30° angular limit from the positions calculated in the 4Xbin refinement. Sub-volumes closer than 120 Å when measured between centers were removed. Sub-volumes with low cross-correlation scores were also removed according to EMAN2’s default settings. Remaining particles were then re-extracted at the original, un-binned pixel size and re-refined starting from previously determined orientations at 2X binning. Sub-tilt refinement in EMAN2 was then carried out using default parameters following sub-tomogram refinement with un-binned sub-volumes (Chen, Bell et al. 2019). For sub-tilt refinement, a threshold mask was used that enclosed only the Env ectodomain portion without any membrane density. The threshold mask was generated using the mask creation tool in the Relion package (Scheres 2012). The final masked ectodomain map contained 32802 sub-volumes with a calculated resolution of 9.13 Å at 0.143 FSC cut-off value.

#### For asymmetric hVLP-Env map

The initial sub-volumes used for generation of the asymmetric map were the same as that used for the C3 symmetrized map described above. Refinement strategies were also nearly identical between the two structures except the asymmetric map was generated with C1 symmetry. However, the refinement and processing steps were carried out completely independent of each other. The number of sub-volumes in the final C1 hVLP-Env structure was 29074 with a global resolution of 10.67 Å at 0.143 FSC cut-off.

#### Structure of hVLP-Env and Gag layer from immature virions

A total of 1520 sub-volumes from only immature virions were used to generate C3 and C1 symmetrized maps of membrane bound Env using 2Xbinned data in EMAN2 (Chen, Bell et al. 2019) by similar procedures as described above.

In the unmasked maps, a third density layer was observed underneath the membrane bilayer. Relaxing symmetry of the C3-density map to C1, showed a more defined organization in the Gag layer. Hence, a short, cylindrical mask enclosing only the Gag layer was used for local refinement of the Gag layer using C1 symmetry, starting from the final refined positions derived from the C3-symmetrized immature Env map. This focused refinement gave rise to a 23 Å map of the Gag protein layer (0.143 FSC cut-off). The refined Gag-CA density map was fitted back into the Gag-CA layer of the relaxed to C1 symmetry full Env map for further analyses.

#### Sub-tomogram averaged structure of BG505-Env from VLPs

Purified BG505-VLPs were mixed with 10nm gold beads (Aurion BSA Gold Tracer 10nm) at a ratio of 15:1 (v/v). The mixture was applied to C-Flat grids and plunge frozen using a Vitrobot Mark IV (FEI Co.) similar to the procedure used for hVLPs above.

Frozen grids were imaged using a 300 kV Titan Krios with a Gatan K2 direct electron detector and GIF energy filter with slit width of 20 eV. Tilt-series were collected in a dose-symmetric tilting scheme from −54° to +54° with a step size of 3° using Leginon (Carragher, Kisseberth et al. 2000) software. Tilt-series were collected in counting mode at a magnification of 53000X, corresponding to a pixel size of 2.58 Å per pixel. The total dose per tilt series was ∼64-68 e^-^/Å^2^. A total of 49 tilt-series were collected.

Tilt-series were imported into EMAN2’s sub-tomogram averaging pipeline (Chen, Bell et al. 2019). 1k X 1k tomograms were generated within EMAN2 using default parameters. Particle points were picked manually in the e2spt_boxer.py interface (Chen, Bell et al. 2019). Further refinement and processing steps were carried out similar to the hVLP-Env sub-tomogram averaging procedures described above. Briefly, sub-volumes were initially extracted at 4xbinning corresponding to a pixel size of 10.32 Å per pixel. Sub-tomogram refinement was carried using a spherical mask including Env on surface and a part of the membrane. The initial model generated for hVLP-Env in Relion (Scheres 2012, Bharat and Scheres 2016) was used as initial model for BG505-Env also with low pass filtering to 60 Å. Sub-volumes were then re-extracted at 2X binning using the particle orientations from the 4X binned refinement output. Sub-tomogram refinement was repeated using the 2X binned data with a cylindrical mask covering only the ectodomain and outer membrane layer. Particle orientations were locally refined, and duplicates were removed. Sub-tilt refinement in EMAN2 was carried out using default parameters (Chen, Bell et al. 2019). The final map contained 2773 sub-volumes with a calculated resolution of 16.7 Å at 0.143 FSC cut-off value.

#### Model fitting and glycan modeling

Full length Env structure from strain 92UG037.8 purified with detergents (PDB ID: 6ULC) (Pan, Peng et al. 2020) was fitted as a rigid body into the 9.13 Å resolution hVLP-Env map. Each protomer, comprising of gp120-gp41 heterodimer, was fitted individually into the map using UCSF Chimera (Pettersen, Goddard et al. 2004). The unstructured loop region at the end of gp41, comprising of amino acid residues 654-664, was fitted into the clearly delineated stalk density using ‘Flexible fitting’ feature in Coot software (Emsley, Lohkamp et al. 2010). For modelling glycan moieties, previously published high-resolution Env structures, PDB IDs: 5FUU and 5FYJ-L, containing well-built glycan models were used as reference (Lee, Ozorowski et al. 2016, Stewart-Jones, Soto et al. 2016). Appropriate glycan chains from these structures were copied into corresponding glycan positions in the hVLP-Env map based on the presence of unoccupied density. In regions where the unoccupied density was larger than the modeled glycan chains available from the reference PDB structures, these extra densities were left un-modeled.

The final model of hVLP-Env, as generated above, was used for rigid body fitting into the other sub-tomogram averaged maps of asymmetric hVLP-Env and immature hVLP-Env. All rigid body fitting procedures were carried out using UCSF Chimera (Pettersen, Goddard et al. 2004).

Root mean square deviation (R.m.s.d) calculations between pairs of HIV Env structures was carried out using UCSF Chimera (Pettersen, Goddard et al. 2004). Multiple sequence alignments were carried out using the Clustal Omega web service (Madeira, Park et al. 2019).

Structure and tomographic images were generated using UCSF Chimera (Pettersen, Goddard et al. 2004), ChimeraX (Pettersen, Goddard et al. 2021), IMOD’s graphical user interface (Kremer, Mastronarde et al. 1996) and ImageJ (Schneider, Rasband et al. 2012).

#### Neutralization assays

ADA.CM.v4 hVLPs were prepared for neutralization assays as described above except omitting the AT-2 inactivation step. Pseudotyped viruses were similarly produced by co-transfection of HEK 293T cells using pSG3ΔEnv backbone plasmid and Env-complementation plasmid. Serial dilutions of antibodies were added to virus and the mixture was incubated for 1 h at 37°C prior to addition to TZM-bl target cells. DEAE-dextran was added to wells to a final concentration of 10 µg/ml. After incubating for 72 h at 37°C, cells were lysed, Bright-Glo luciferase reagent (Promega) was added, and luminescence was measured using a Synergy H1 microplate reader (Bio-Tek).

#### Virus ELISA

ELISAs on directly immobilized hVLPs were performed as previously described (Tong, Crooks et al. 2012). Briefly ADA.CM.v4 hVLPs (prepared as detailed above and treated with or without AT-2) were immobilized on microwell plates at a 20x concentration for 2 h at 37°C. Plates were blocked with 4% non-fat dry milk (NFDM) PBS for 1 h at 37°C, probed with serial dilutions of primary antibodies and, subsequently, goat anti-human-Fcɣ-HRP secondary antibody (Jackson) in PBS + 0.4% NFDM, and with washes using PBS between each step (detergent was omitted from all steps). Signal was developed using One-step Ultra TMB Substrate (ThermoFisher).

#### Light microscope imaging of infected cells

TZM-bl target cells were plated in 96-well plates (100 µl at 1 x 10^5^/ml), and 24 hours later an equal volume of hVLPs were overlaid on the cells. After a 72 hour incubation, cells were imaged using a Evos FL light microscope (Life Technologies) and 20x objective.

#### BG505.SOSIP expression and purification

Soluble BG505.SOSIP protein was expressed and purified as previously described (Verkerke, Williams et al. 2016). Expi293F cells grown in Freestyle 293 media (Thermo Scientific) were co-transfected with plasmids containing the SOSIP sequence and furin protease in a 3:1 ratio using PEI MAX (Polysciences). Six to seven days post-transfection the supernatant was cleared by filtration and the SOSIP purified by *Galanthus nivalis* lectin affinity, followed by ion-exchange and hydrophobic interaction chromatography. Trimeric product was buffer exchanged into PBS and stored at 4°C.

### Hydrogen-deuterium mass-spectroscopy (HDX-MS) of ADA.CM.v4 particles and BG505.SOSIPs

#### HDX-MS using quench lysis method

Frozen stocks of ADA.CM.v4 VLP containing approximately 0.5mg/mL envelope protein or purified BG505.SOSIP diluted to the same concentration were buffer-exchanged into HEPES-buffered saline (HBS, 10mM HEPES-KOH, pH 7.4, 150mM NaCl) using Zeba Spin columns with a 7kDa MWCO (Thermo Fisher). 10µL of VLP solution was diluted with 90µL of deuterated HBS (Hepes Buffer Saline, pH 7.4) for the indicated labeling time. The reaction was quenched and lysed by addition of an equal volume of 0.2% formic acid, 4M guanidine hydrochloride, 0.2M TCEP (tris(2-carboxyethyl)phosphine), 0.2% DDM (n-Dodecyl-B-D-maltoside), followed by 30s incubation on ice. 20µL of a 300mg/mL solution of ZrO_2_ beads (HybridSPE-Phospholipid) (Adhikary, Deredge et al. 2017) in 0.1% formic acid was then added to the solution, followed by vortexing for 30seconds on ice. The bead/protein mixture was then moved to a 0.22µm cellulose acetate centrifuge filter tube (Spin-X, Corning) and spun for 30s at 13,000xG, 0°C. The flow-through was transferred to a thin-wall PCR tube and frozen in liquid nitrogen. Samples were thawed on ice and passed over a custom packed pepsin column (2.1 × 50 mm) kept at 15°C with a flow of 0.1% trifluoroacetic acid (TFA), 2% acetonitrile (ACN) at 200 μL/min for 5 minutes.

Digested peptides were collected on a Waters XSelect CSH C18 XP VanGuard Cartridge, 130Å, 2.5 µm, 2.1 mm X 5 mm before separation on a Waters ACQUITY UPLC Peptide CSH C18 Column, 130Å, 1.7 µm, 1 mm X 100 mm using a gradient of 7 to 14% B over 1 minute, 14 to 30% B over 12.5 minutes, and 30 to 50% B over 1 minute, followed by washing with three rapid gradients between 95 and 5% B (A: 0.1% formic acid, 0.025% trifluoroacetic acid, 2% acetonitrile; B: 0.1% formic acid in 100% acetonitrile). The liquid chromatography system was coupled to a Waters Synapt G2-Si Q-TOF with ion mobility enabled. Source and de-solvation temperatures were 70°C and 130°C respectively. The StepWave ion guide settings were tuned to prevent non-uniform gas phase proton exchange in the source (Guttman, Wales et al. 2016).

During the separation step, a series of 250 μL injections were used to clean the pepsin column: (1) Fos-choline-12 (Anaspec) in 0.1% TFA; (2) 2 M GuHCl in 0.1% TFA; (3) 10% acetic acid, 10% ACN and 5% IPA (Majumdar, Manikwar et al. 2012, Hamuro and Coales 2018). After each gradient, the trap column was washed with a series of 250 μL injections: (1) 10% FA; (2) 30% trifluoroethanol; (3) 80% MeOH; (4) 66% 2-propanol, 34% ACN; (5) 80% ACN (Fang, Rand et al. 2011).

#### HDX-MS using solution digestion method

Env on hVLPs was quantified by SDS-PAGE using BG505 SOSIPs as a standard. HDX-MS reactions were initiated by diluting 20 uL of either hVLPs or BG505 SOSIPs with 180 uL deuteration buffer (10 mM Phosphate, 150 mM NaCl, 85% D_2_O (Cambridge Isotope Laboratories)) to a final pH = 7.45. Samples were deuterated for 5 seconds, 60 seconds, 30 minutes, or 3 hours before being diluted 1:1 with ice cold quench buffer (8 M urea, 200 mM TCEP [tris(2-carboxyethyl) phosphine] and 0.2% formic acid (FA)) to a final pH of 2.5. Quenched samples were digested with 30 ug/mL of porcine pepsin (Worthington Labs) under quench conditions for 5 minutes on ice. Labeled peptides were purified by high-speed centrifugation at 0°C (2 minutes at 25,000 rcf) and immediately flash frozen in liquid nitrogen. BG505 SOSIP samples were handled identically to ensure consistent labeling and back exchange. Frozen samples were stored at −80°C until analysis. All reactions were performed in duplicate.

Samples were thawed for 5 minutes on ice and manually injected into a custom built HDX LC system kept at 0°C using a 500 uL sample loop. Samples were trapped on a Waters ACQUITY UPLC CSH C18 VanGuard 130Å, 1.7 μm, 2.1 mm by 5 mm trap column for 7 minutes with a flow of solvent A [2% acetonitrile, 0.1% FA, 0.025% trifluoroacetic acid (TFA)] at a rate of 150 μL/min. Peptides were resolved over a Waters ACQUITY UPLC CSH C18 130Å, 1.7 μm, 1 x 100 mm column using a 20 minute linear gradient of 3% to 50% solvent B (Solvent B: 100% acetonitrile and 0.1% FA) and analyzed using Waters Synapt G2-Si Q-TOF as described above. Following each injection, the sample loop and trap were washed as described above.

Deuterium uptake analysis was performed with HD-Examiner (Sierra Analytics) and HX-Express v2 (10.1021/jasms.8b04435). For solution digestion experiments data was extracted using CDCReader and analyzed using HXExpress V2 (Guttman, Weis et al. 2013) with binomial fitting and bimodal deconvolution.

Internal exchange standards Pro-Pro-Pro-Ile [PPPI] and Pro-Pro-Pro-Phe [PPPF] were included in each reaction to control for variations in ambient temperature during the labeling reactions (10.1021/ac300535r). Back-exchange was measured by including bradykinin peptide at 2µg/mL in the deuterated buffer to serve as a fully deuterated control. The back-exchange level ranged from 12-16% across experiments; deuterium uptake was not corrected.

Totally deuterated (TD) samples were prepared by collecting purified peptide eluent following reverse phase LC separation of a pepsin digested undeuterated sample. Following evaporation of the LC elution buffer the peptides were resuspended in HDX PBS pH 7.50 Buffer, deuterated in deuteration Buffer for 1 hours at 65°C, and quenched and frozen as described above.

For peptide identification, un-deuterated peptides were collected from the LC system, dried by speed-vac, and resuspended in 5% acetonitrile, 0.1% formic acid for re-injection on an Orbitrap Fusion for MS/MS using EThcD fragmentation. Data was analyzed using Byonic (Protein Metrics) and manually compared to undeuterated sample data.

#### Peptic digest analysis

Peptic digest products were collected from the HDX chromatography system, dried by speed-vacuum, and resuspended in aqueous buffer for nanoscale liquid chromatography (nanoLC-MS) using a 90-minute linear gradient from 2-40% acetonitrile. Products were analyzed on an Orbitrap Fusion mass spectrometer (ThermoFisher Scientific) using a high-energy collisional dissociation (HCD) product-dependent electron-transfer/high-energy collision dissociation (EThcD) method with a targeted mass list method with a targeted mass list using HexNAc, HexHexNAc, and Hex2HexNAc m/z (204, 366, and 528 respectively) to trigger EThcD. Glycopeptide data were visualized and processed by Byonic™ (Version 3.8, Protein Metrics Inc.) using a 10 ppm precursor and 10 ppm fragment mass tolerance. Glycopeptides were searched using the N-glycan 309 mammalian database in Protein Metrics PMI-Suite and scored based on the assignment of correct c- and z-fragment ions. The true-positive entities were further validated by the presence of glycan oxonium ions m/z at 204 (HexNAc ions) and 366 (HexNAcHex ions).

## Supplemental Information

Supplemental information can be found online at ***

## Supplemental Figures

**Figure S1.**
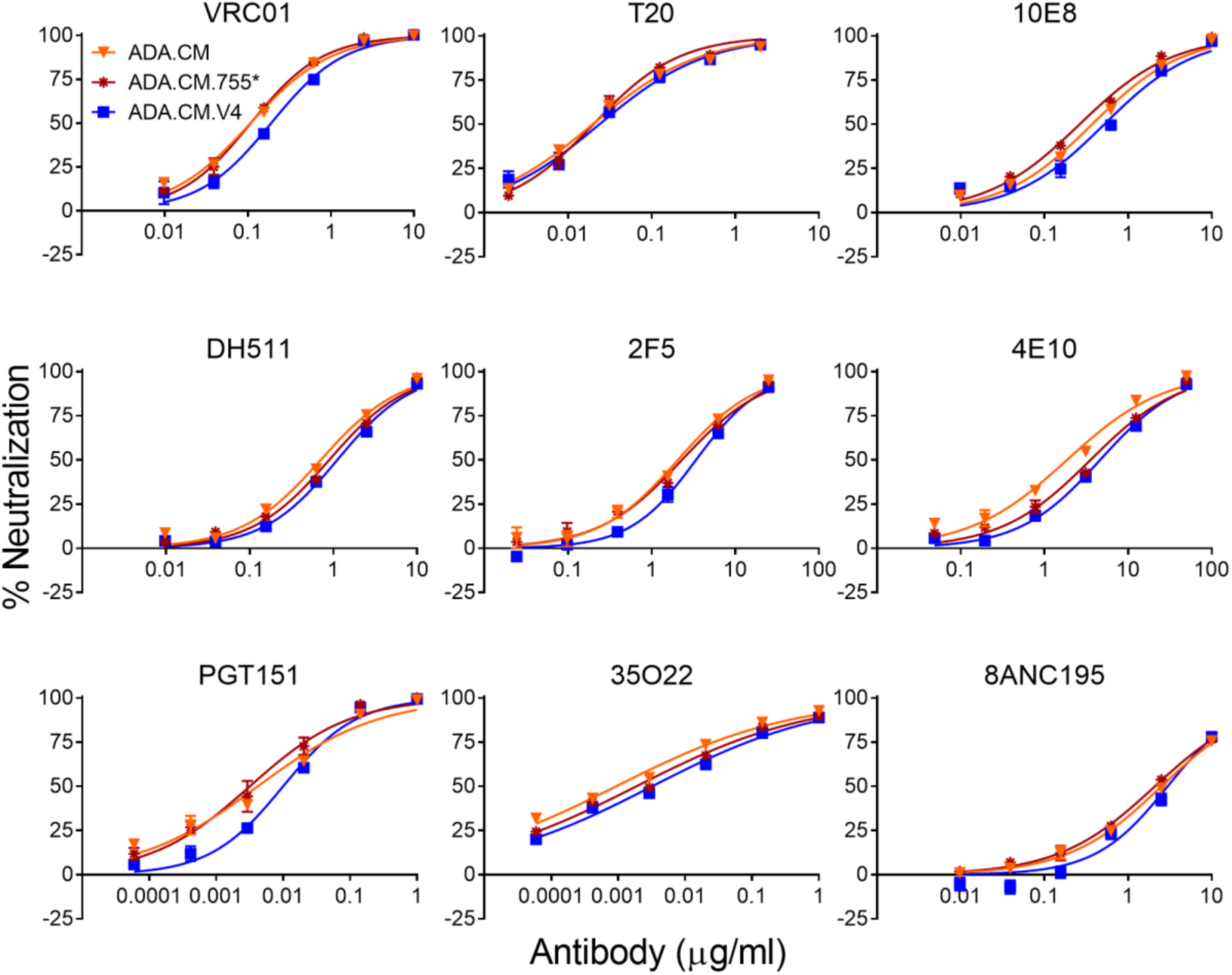
Percentage neutralization of HIV particles using a multi-antibody panel and the gp41 fusion inhibitor peptide, T20. Related to Figure 1. ADA.CM and ADA.CM.755* denote pseudotyped HIV particles produced by transient transfection of HEK293T cells using a complementary two-plasmid method, with one plasmid DNA bearing the HIV backbone genes except Env, and the other plasmid bearing either Env displaying full length ADA.CM Env, or ADA.CM with a CTD truncation similar to that in hVLPs (stop codon at amino acid 756), respectively. ADA.CM.V4 refer to hVLPs produced by transient transfection of the ADA.CM.V4 high-Env HEK293T stable cell line with plasmid DNA bearing the HIV backbone genes except Env.

**Figure S2.**
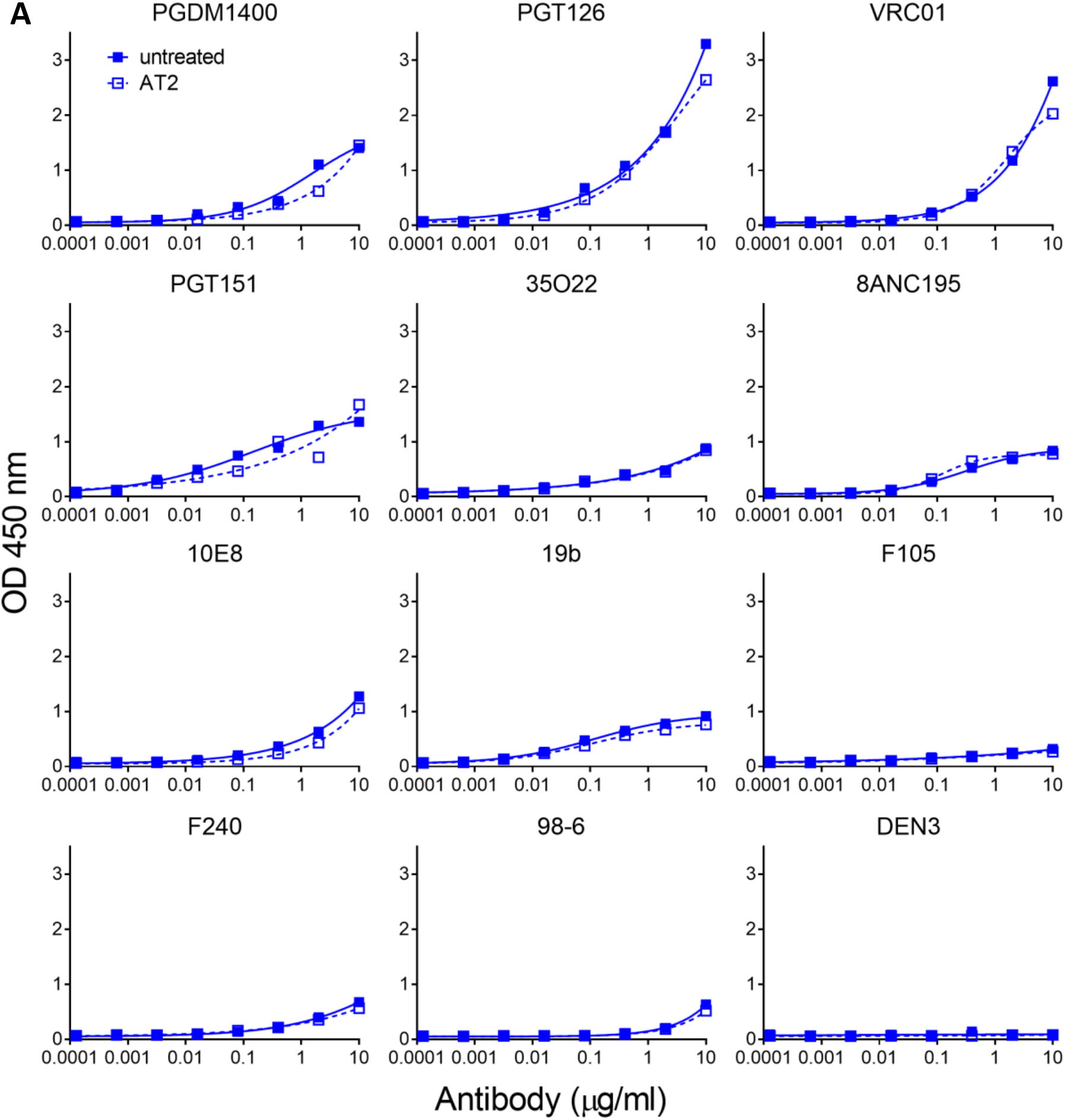
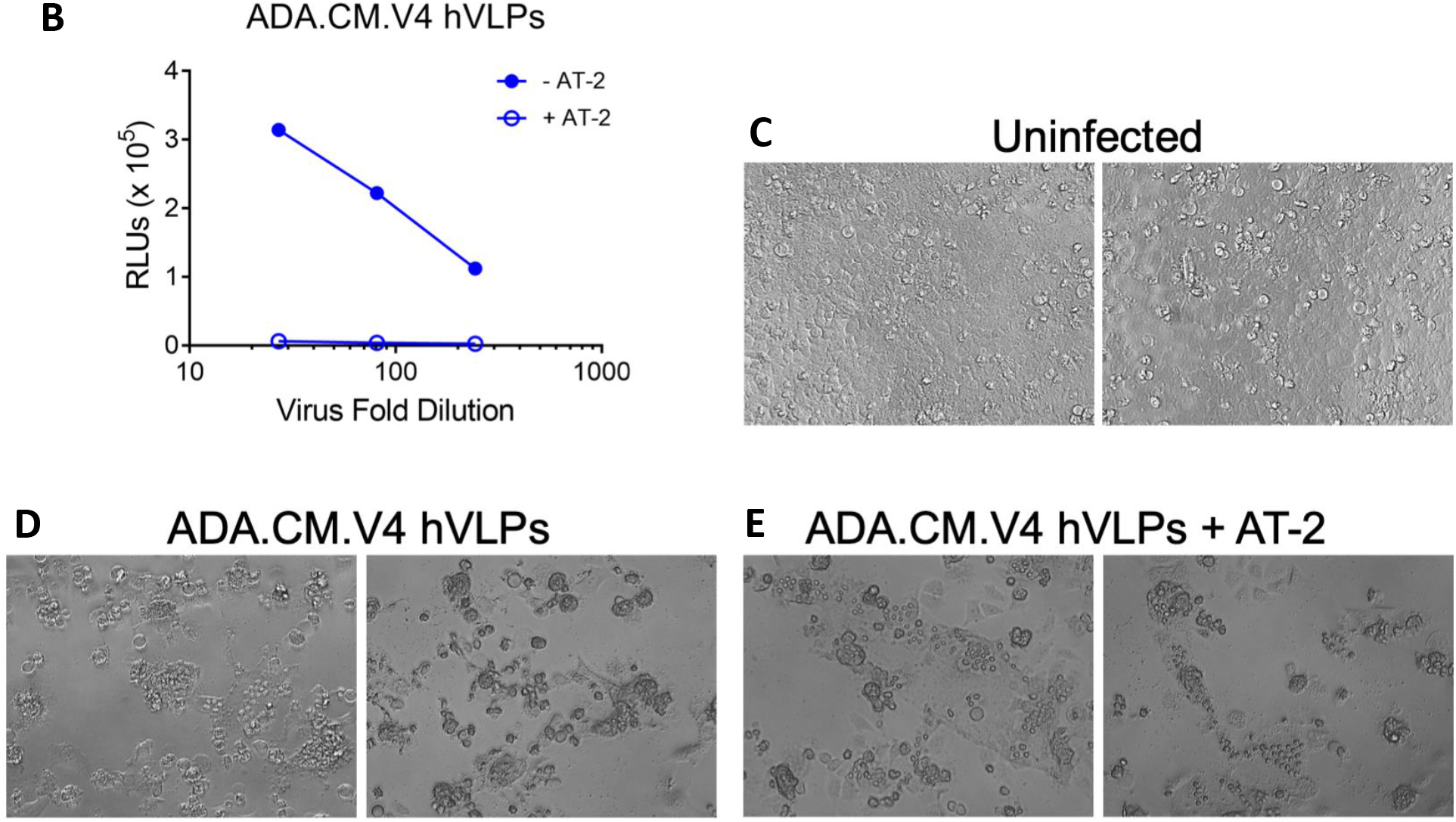
Effect of AT-2 treatment on hVLPs. Related to Figure 1. **(A)** Binding of antibodies probed to hVLPs, as probed using an ELISA. hVLPs were treated in presence (open squares) or absence (closed squares) of AT-2 and directly immobilized on microwell plates. Binding curves show no significant difference in antibody binding on treatment with AT-2, indicating that the antigenic surface of hVLP-Env has not been altered due to AT-2 inactivation. **(B)** ADA.CM.V4 hVLPs, treated in presence (open circles) and absence (close circles) of AT-2 and normalized for p24 content, were overlaid on TZM-bl reporter cells and luciferase expression was monitored 72 hours later. AT-2-treated hVLPs do not cause productive infection. **(C-E)** Cells that had been incubated with hVLPs as in **(B)** were imaged using a light microscope and representative images are shown. Uninfected cells are monodispersed, cells incubated with hVLPs (treated either in presence or absence of AT-2) induce syncytia formation and cytotoxicity. Images are from wells in which hVLP containing supernatants were effectively diluted 3-fold prior to addition to cells.

**Figure S3.**
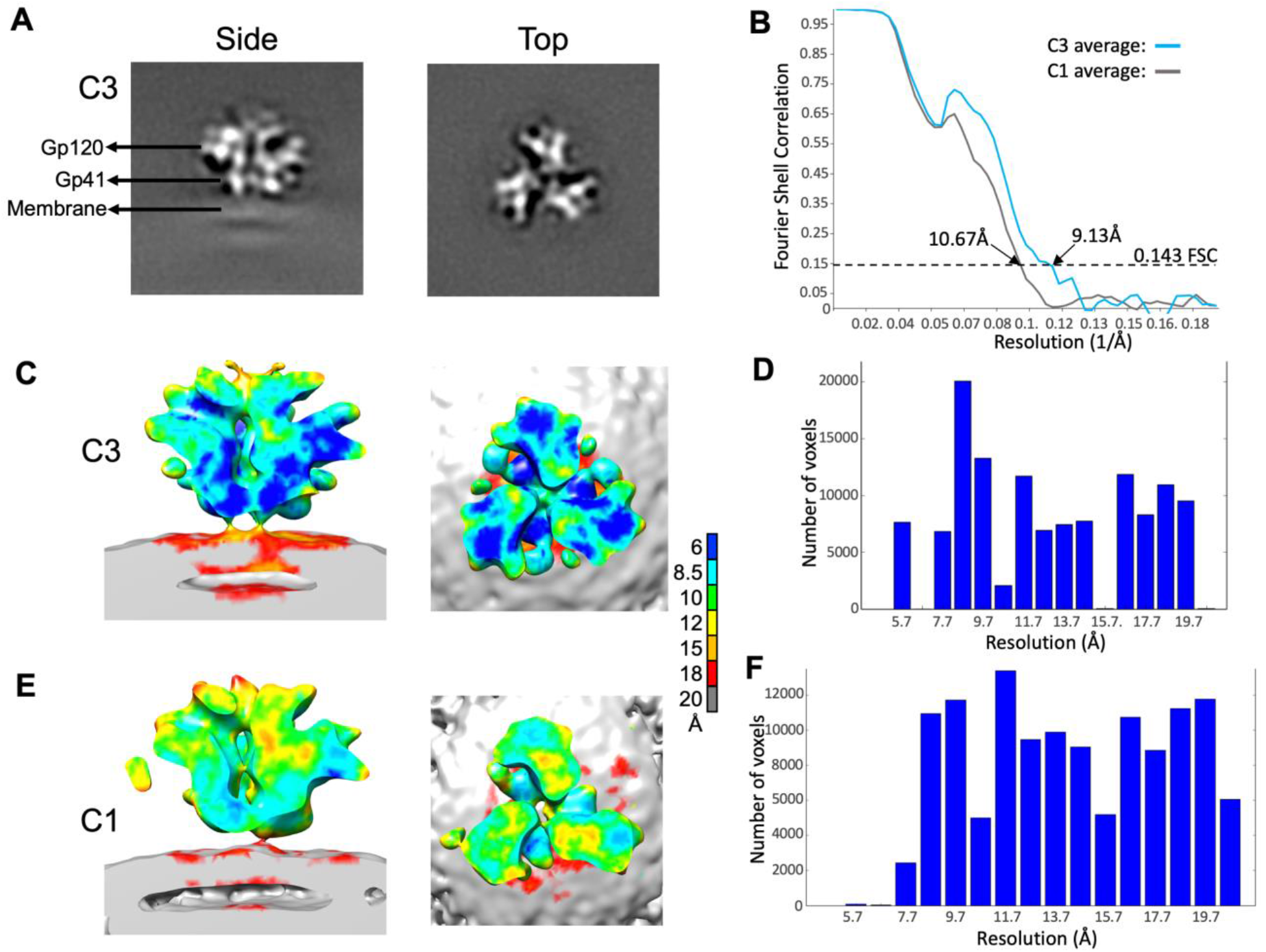
Resolution estimates for hVLP-Env sub-tomogram averaged maps. Related to Figures 1, 3 and 5. **(A)** Perpendicular slices through the C3-symmetrized hVLP-Env map. **(B)** Gold-standard Fourier shell correlation (FSC) curves for C3-symmetrized map (blue line) and C1-symmetry map (grey line). Dotted line indicated the 0.143 FSC cutoff. **(C-F)** Local resolution estimation using Resmap software (Kucukelbir, Sigworth et al. 2014) for C3-symmetrized map (C,D) and asymmetric map (E,F).

**Figure S4.**
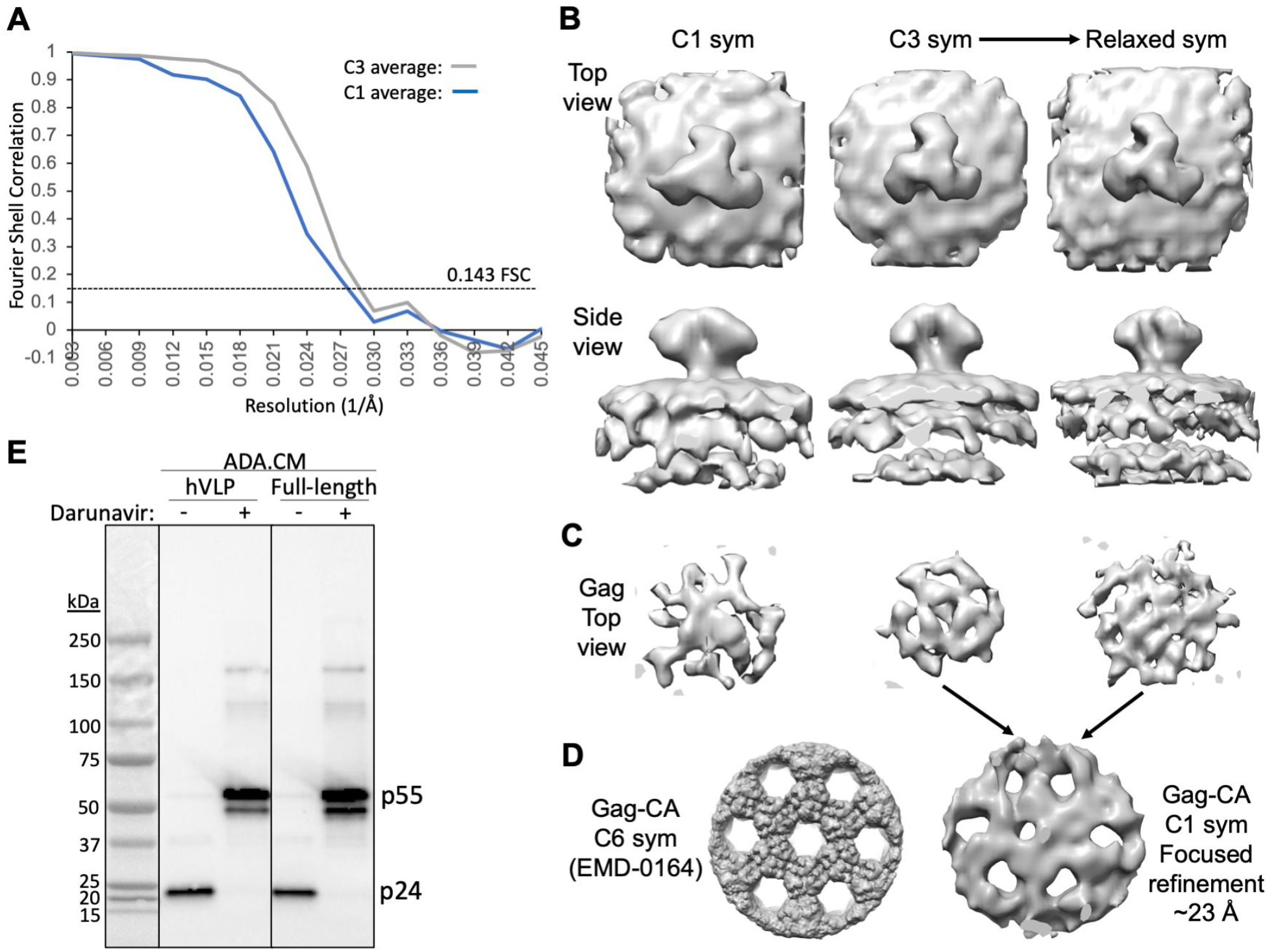
Structural analysis of Env, Gag-MA and Gag-CA proteins in immature hVLPs. Related to Figure 1 and 2. **(A)** Fourier shell correlation (FSC) curves for C3 (grey line) and C1 (blue line) symmetrized maps of immature hVLP-Env. **(B)** Top and side surface views of immature hVLP-Env structure with C1 symmetry, C3 symmetry and the C3 map relaxed to C1 symmetry. **(C)** Top view of Gag layer from corresponding averages in panel B. **(D)** Left: Previously published high resolution density map of Gag-CA lattice, low pass filtered to ∼10 Å (Mattei, Tan et al. 2018). Right: Gag-CA structure derived after focused refinement of Gag layer from the relaxed C1 symmetry immature Env map. **(E)** HIV protease inhibitor darunavir blocks the processing of immature pr55 Gag precursor into the mature p24 capsid protein. ADA.CM Env with 102 residues truncation in CTD from hVLPs and full-length Env-CTD from pseudovirus, produced in the presence (+) or absence (-) of darunavir, were run on an SDS-PAGE gel and probed by Western blot staining with an anti-p24 antibody.

**Figure S5.**
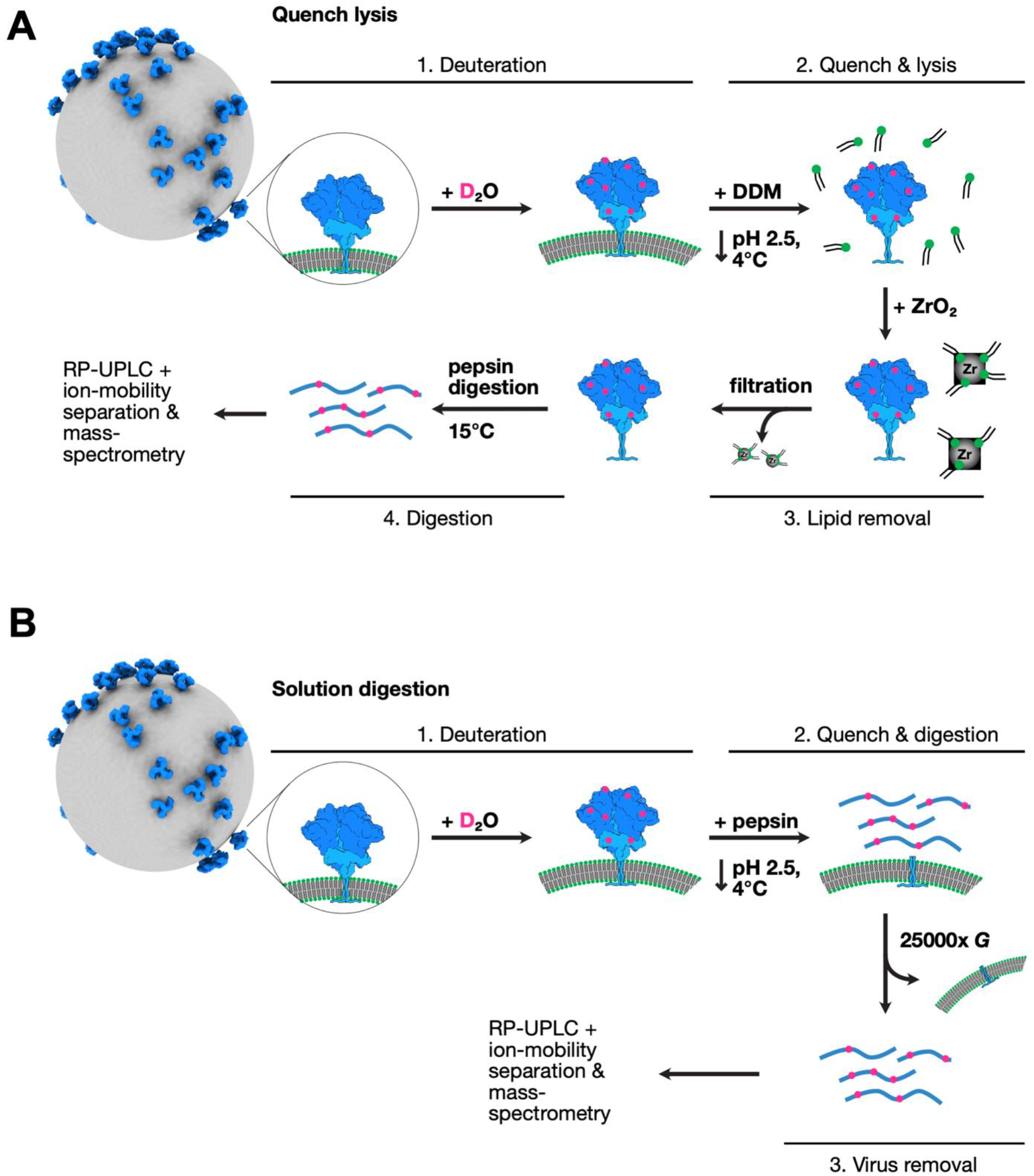
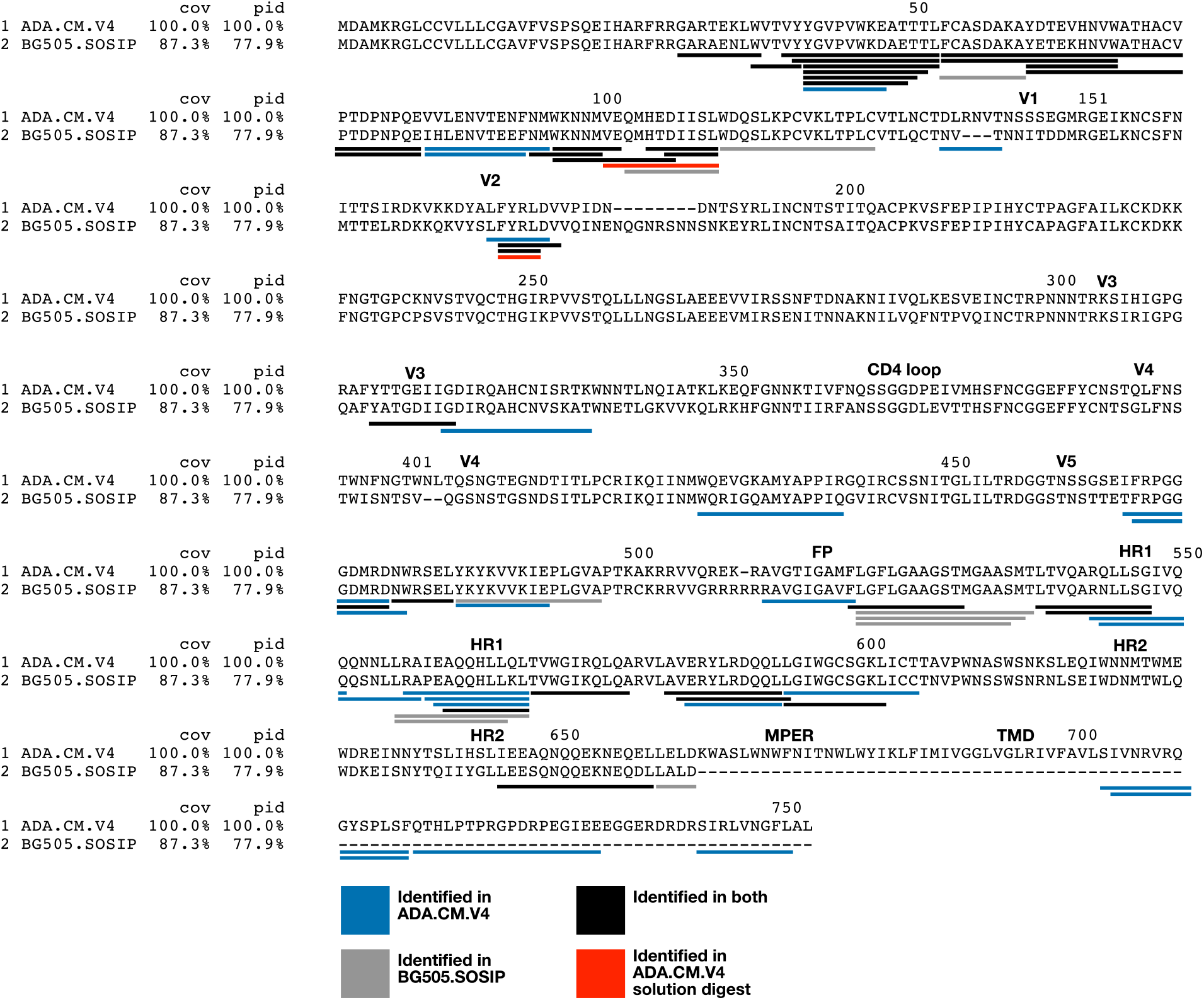
Hydrogen-deuterium exchange mass spectroscopy (HDX-MS) methods used for hVLP-Env analysis. Related to Figures 3 and 5. **(A)** Quench Lysis method: Improved procedure to increase peptide coverage in HDX-MS experiment. hVLP particles were treated with deuterated water (D_2_O) in native, neutral pH conditions for varying periods of time. The reaction was quenched along with detergent treatment to solubilize Env. Zirconium beads (Zr) were used to filter out the lipid molecules before downstream peptidase digestion and mass-spectrometry analysis (see Methods for details). **(B)** Solution digestion method: Routine HDX-MS procedure where after deuteration of hVLP-Env in native, neutral pH, the reaction is quenched, and peptidase treatment is carried out with Env fixed on particle surface. **(C)** Peptide coverage map of hVLP-Env by quench lysis HDX-MS. Sequence alignment of ADA.CM and BG505.SOSIP used in the HDX-MS experiments. BG505.SOSIP includes residues only upto 664 and does not contain MPER, TMD or CTT domains. Underlined peptide regions are color coded based on their identification in the quench lysis HDX-MS experiment. Peptides underlined in red are those identified by solution digest HDX-MS. Important regions of Env such as variable loops (V1-V5), CD4 receptor binding loop, fusion peptide (FP), heptad repeat helices (HR1 and HR2), MPER and TMD are highlighted and labeled.

**Figure S6.**
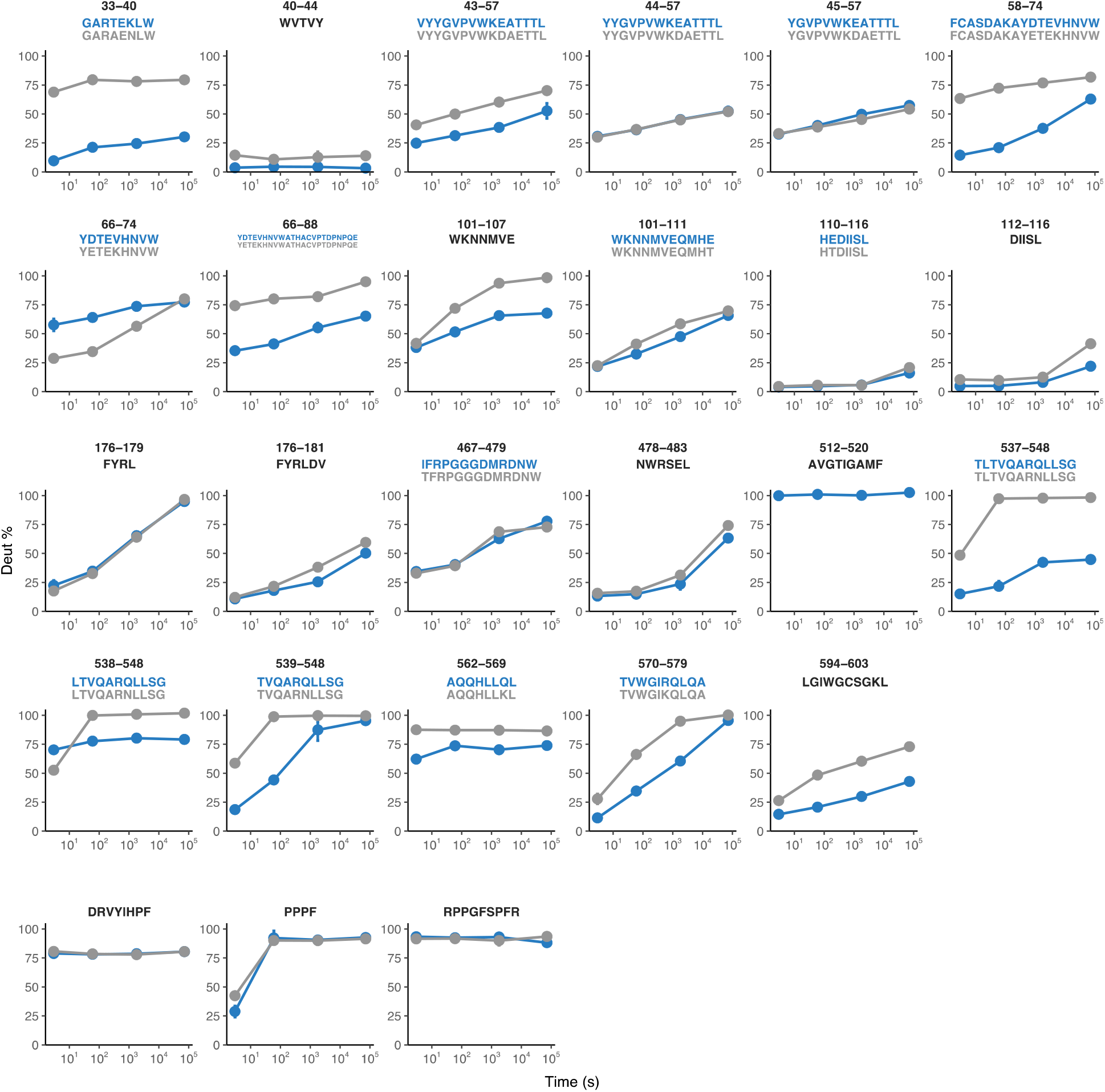
Deuterium uptake plots from HDX-MS experiment. Related to Figures 3 and 5. Deuterium uptake plots for all peptides in Env ectodomain commonly identified between hVLP-Env (blue) and BG505.SOSIP (grey). Corresponding sequence of the peptides are given above each uptake plot. Bottom three panels are back-exchange peptide standards (DRVYIHPF and RPPGFSPFR) and exchange rate peptide standard (PPPF). Refer to Figure S10 for sequence alignment between the two Envs.

**Figure S7.**
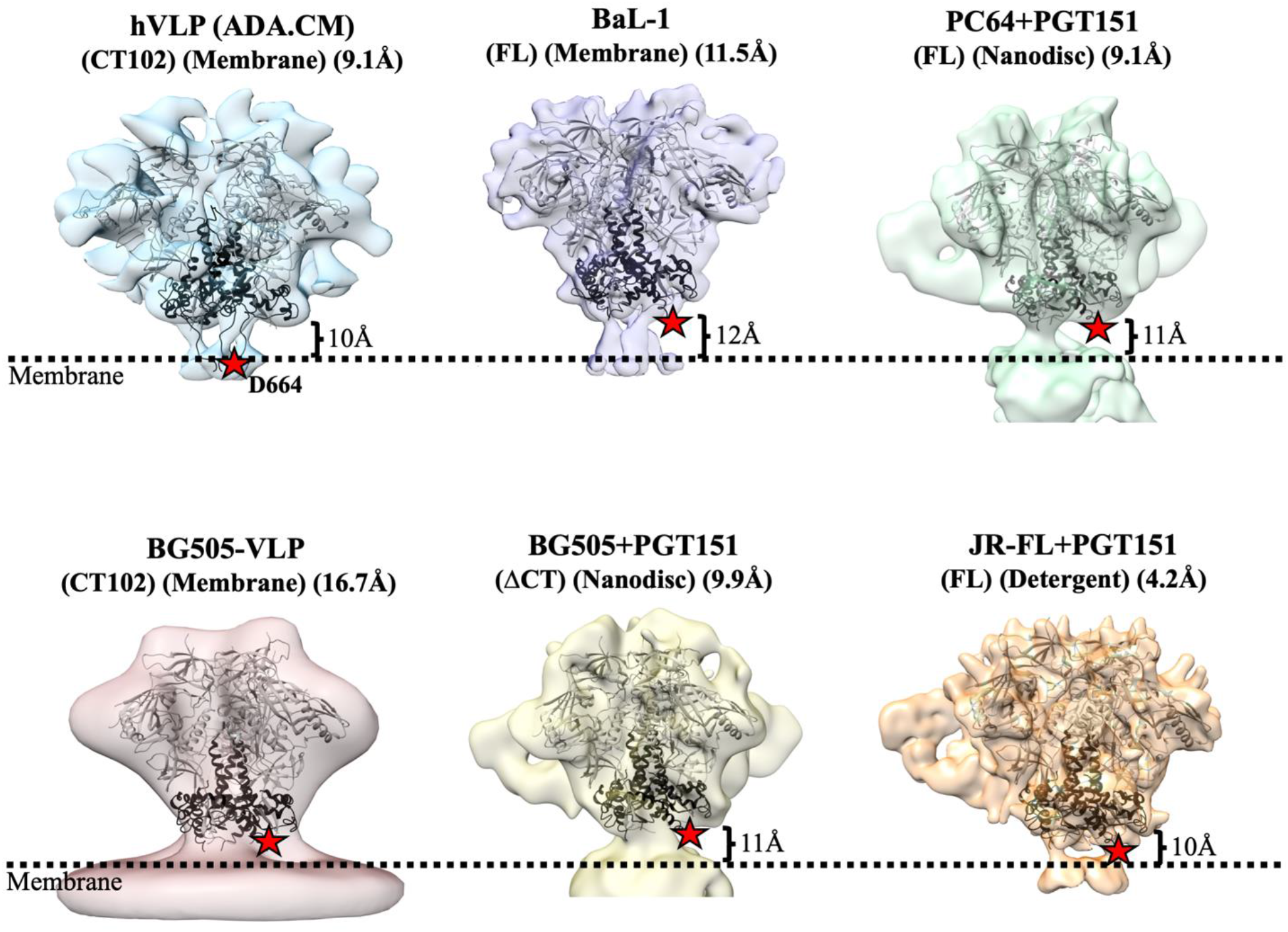
Position of Env ectodomain with respect to membrane surface. Related to Figure 3. Surface view of Env structure from different HIV strains: hVLP (ADA.CM), BaL-1, PC64, BG505 and JR-FL. Position of outer membrane surface as interpreted for each of the structures is indicated by a dotted black line. Position of Asp-664, which forms the end of HR2 helix region in gp41, is denoted by a red star in all structures. Low resolution structure of near full length BG505 in native membrane was obtained from VLPs displaying high levels of BG505 Env and having a CTD truncation similar to that in hVLP-Env.

**Figure S8.**
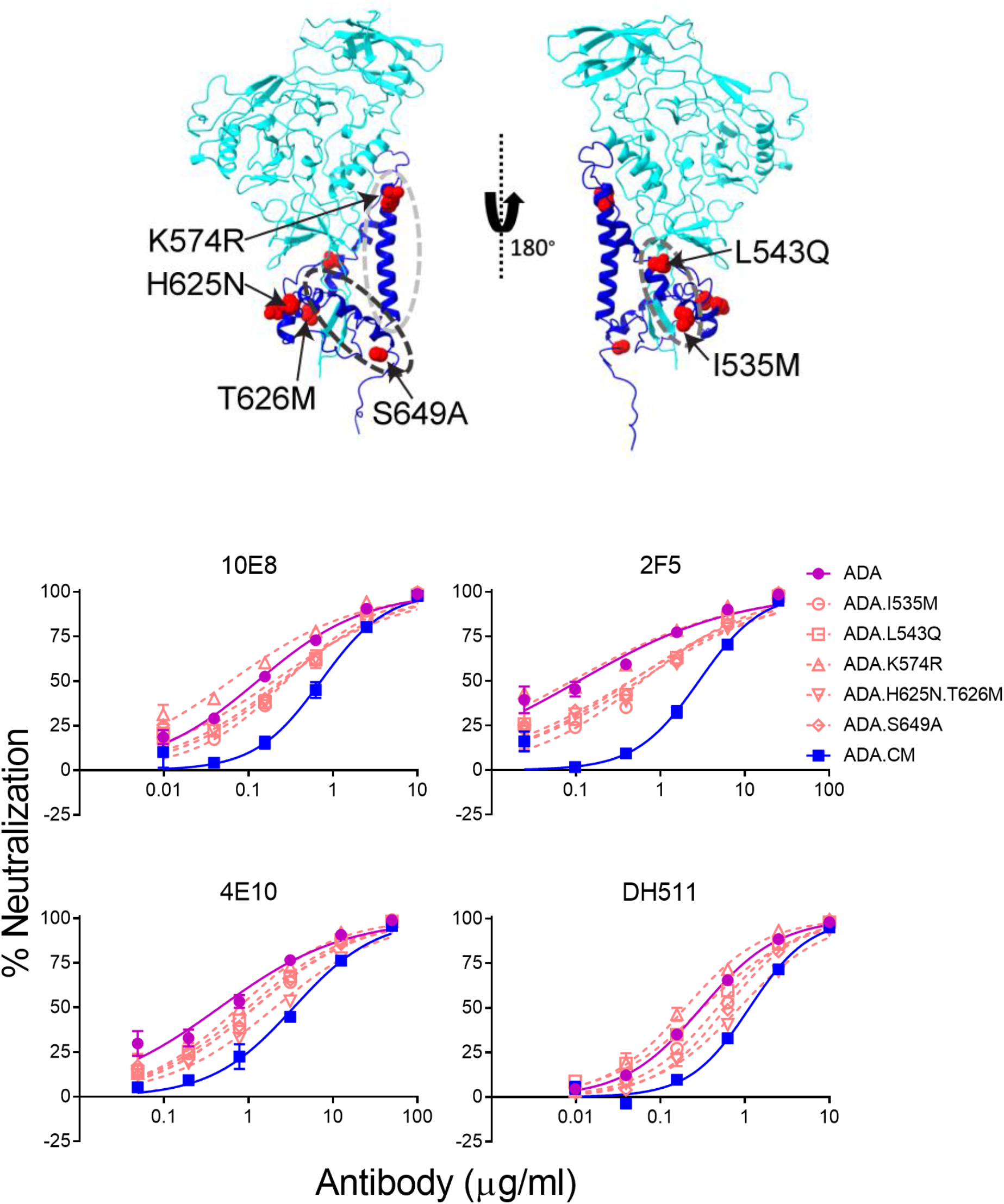
Neutralization effect of individual mutations in gp41 subunit of ADA.CM with respect to MPER-targeting antibodies. Related to Figure 4. Top: Mutations relative to parental ADA virus in gp41 subunit that contribute to ADA.CM trimer hyperstability mapped onto its structure. The mutated residues are represented as red balls. HR1-C helix, HR2 helix and fusion loop proximal region are indicated by light grey, black and dark grey ovals respectively. Bottom: MPER antibody neutralization of single residue mutants of HIV-1 ADA in gp41 subunit, as described in Figure S14. Each ADA point mutant corresponds to residues in ADA.CM that were mutated during directed evolution of ADA to withstand destabilizing conditions (Leaman and Zwick 2013). Curves shown are from a single experiment performed in duplicate.

**Figure S9.**
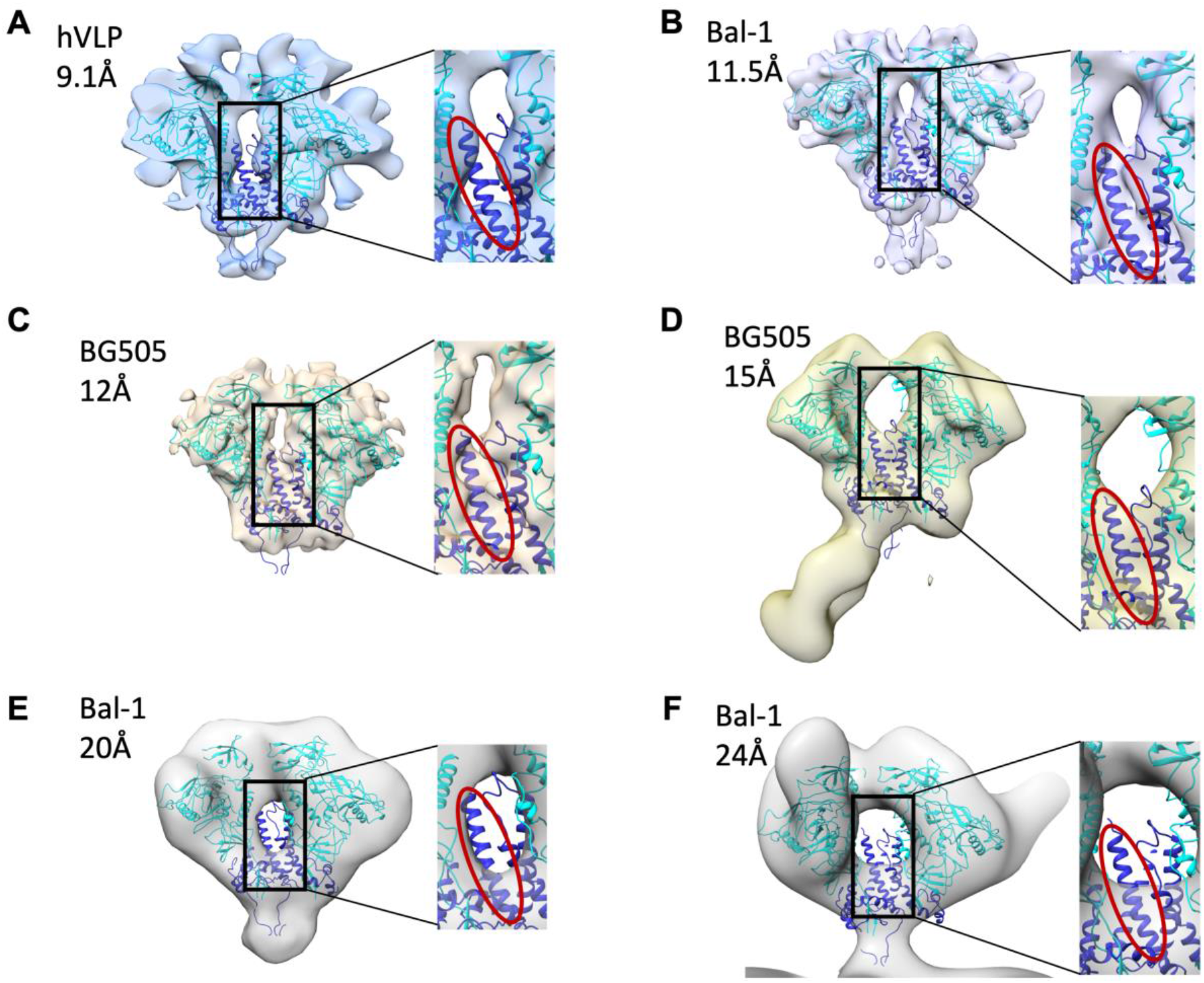
Comparison of HR1-C density in Env trimer structures at different resolutions. Related to Figure 5. **(A-F)**. Surface slab view of Env single particle cryo-EM and sub-tomogram averaged reconstructions with only 2 protomers shown to visualize HR1-C helix density clearly. In each panel, the central core region (black rectangle) is zoomed in and the HR1-C helix indicated with red oval. Maps shown are (A) hVLP-Env, (B) BaL-1 (EMD-21412) (Li, Li et al. 2020), (C) BG505 SOSIP.664 (EMD-5782) (Lyumkis, Julien et al. 2013), (D) BG505 SOSIP v5.2 (EMD-21075) (Cottrell, van Schooten et al. 2020), (E) BaL-1 (EMD-5019) (Liu, Bartesaghi et al. 2008), (F) Bal-1 (EMD-5457) (Tran, Borgnia et al. 2012).

## Supplemental Tables

**Table S1.** Half-maximal inhibitory concentration (IC_50_) values for antibody neutralization of VLPs displaying Envs of full-length ADA.CM, ADA.CM with truncation matching that of hVLPs (ADA.CM.755*) and high Env displaying hVLPs (ADA.CM.V4).

|  |  | ADA.CM | ADA.CM.755* | ADA.CM.V4<br>(hVLP) |
| --- | --- | --- | --- | --- |
| IC<sub>50</sub> (µg/ml) | VRC01 | 0.11 | 0.11 | 0.19 |
|  | T-20 | 0.020 | 0.020 | 0.024 |
|  | 10E8 | 0.37 | 0.26 | 0.52 |
|  | DH511 | 0.71 | 0.89 | 1.12 |
|  | 2F5 | 2.06 | 2.31 | 3.39 |
|  | 4E10 | 1.87 | 3.57 | 4.64 |
|  | PGT151 | 0.0042 | 0.0032 | 0.0099 |
|  | 35O22 | 0.0093 | 0.0019 | 0.0032 |
|  | 8ANC195 | 2.51 | 2.10 | 2.98 |

**Table S2.** IC_50_ values for neutralization of HIV-1 strains by MPER-targeted antibodies.

|  |  | ADA.CM.V4 | BG505 | BaL |
| --- | --- | --- | --- | --- |
|  | MPER-targeting antibodies |  |  |  |
| IC<sub>50</sub> (µg/ml) | 10E8 | 0.46 | 0.16 | 0.21 |
|  | DH511 | 1.26 | 0.21 | 1.00 |
|  | 2F5 | 2.88 | 0.94 | 1.36 |
|  | 4E10 | 5.16 | 0.81 | 1.83 |
|  | Gp120-gp41 interface and glycan targeting antibodies |  |  |  |
|  | VRC01 | 0.16 | 0.0086 | 0.0028 |
|  | PGT151 | 0.011 | 0.00026 | 0.00083 |
|  | 35O22 | 0.0032 | >20 | 0.00018 |
|  | 3BC176 | 66.7 | 36.5 | 0.93 |

**Table S3.** IC_50_ values for neutralization of single point mutants in gp41 of ADA by MPER-targeted antibodies.

|  | IC<sub>50</sub> (μg/ml) |  |  |  |  |  |
| --- | --- | --- | --- | --- | --- | --- |
|  | VRC01 | T20 | 10E8 | DH511 | 2F5 | 4E10 |
| ADA | 0.31 | 0.016 | 0.13 | 0.31 | 0.11 | 0.46 |
| ADA.I535M | 0.36 | 0.013 | 0.28 | 0.47 | 0.60 | 1.17 |
| ADA.L543Q | 0.39 | 0.072 | 0.24 | 0.32 | 0.46 | 0.99 |
| ADA.K574R | 0.27 | 0.024 | 0.061 | 0.20 | 0.089 | 0.76 |
| ADA.H625N.T626M | 0.41 | 0.11 | 0.26 | 0.86 | 0.56 | 2.05 |
| ADA.S649A | 0.29 | 0.021 | 0.20 | 0.60 | 0.44 | 1.18 |
| ADA.CM | 0.16 | 0.013 | 0.72 | 1.13 | 2.86 | 3.43 |

**Table S4.**
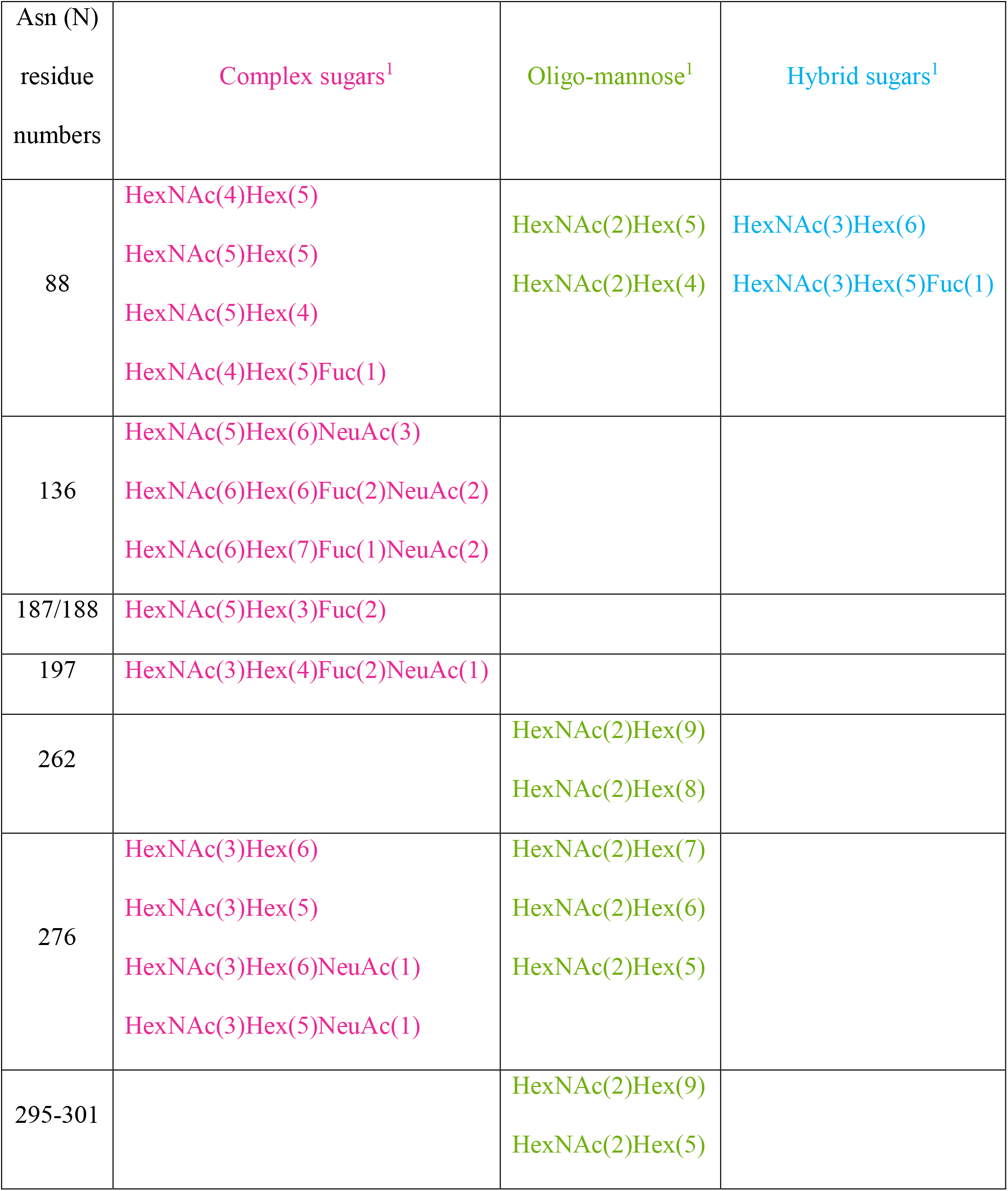
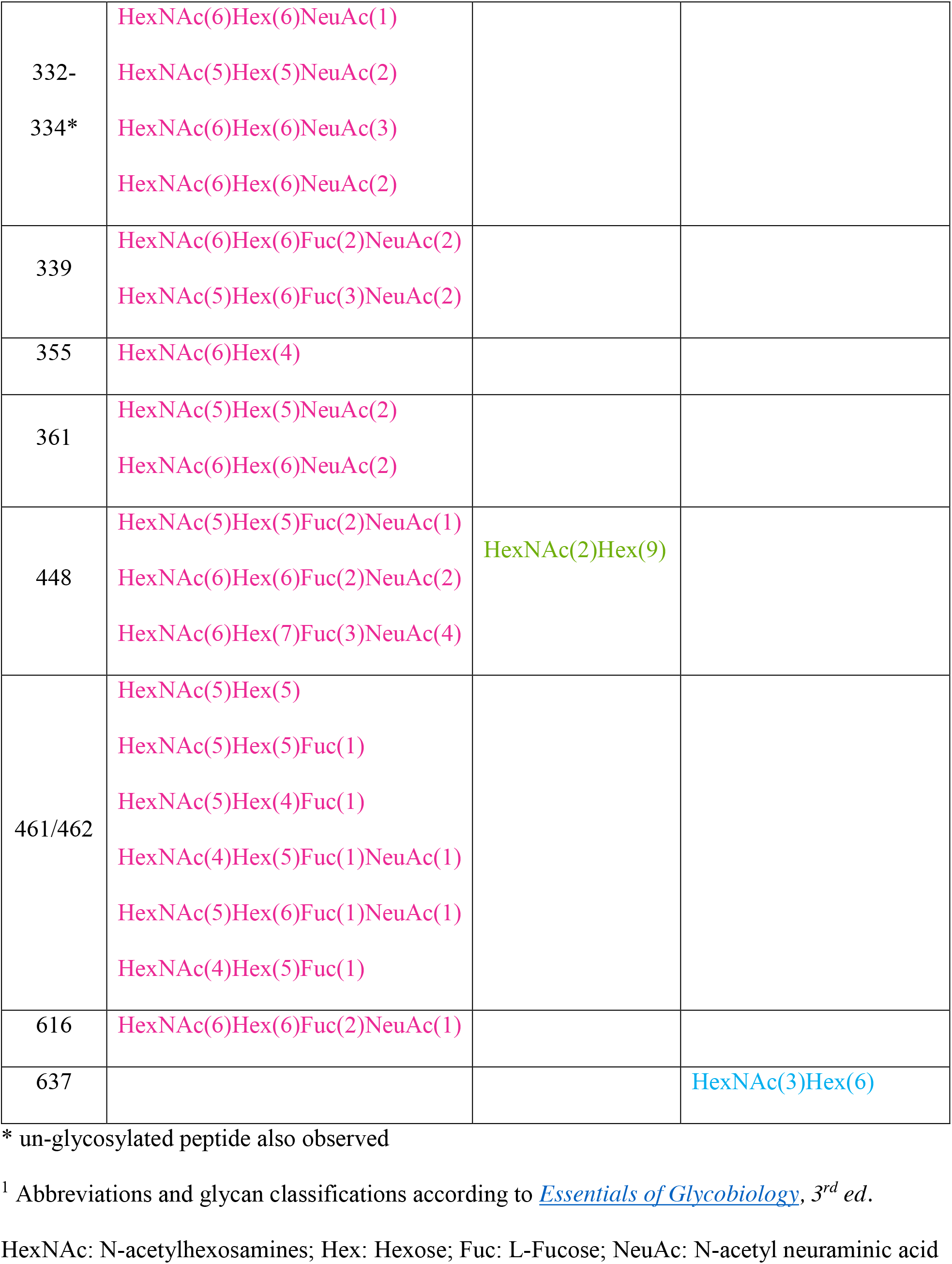
N-linked glycosylation detected in hVLP-Env through peptide digest analysis by mass spectrometry.

| Asn (N)<br>residue<br>numbers | Complex sugars<sup>1</sup> | Oligo-mannose<sup>1</sup> | Hybrid sugars<sup>1</sup> |
| --- | --- | --- | --- |
| 88 | HexNAc(4)Hex(5)<br>HexNAc(5)Hex(5)<br>HexNAc(5)Hex(4)<br>HexNAc(4)Hex(5)Fuc(1) | HexNAc(2)Hex(5)<br>HexNAc(2)Hex(4) | HexNAc(3)Hex(6)<br>HexNAc(3)Hex(5)Fuc(1) |
| 136 | HexNAc(5)Hex(6)NeuAc(3)<br>HexNAc(6)Hex(6)Fuc(2)NeuAc(2)<br>HexNAc(6)Hex(7)Fuc(1)NeuAc(2) |  |  |
| 187/188 | HexNAc(5)Hex(3)Fuc(2) |  |  |
| 197 | HexNAc(3)Hex(4)Fuc(2)NeuAc(1) |  |  |
| 262 |  | HexNAc(2)Hex(9)<br>HexNAc(2)Hex(8) |  |
| 276 | HexNAc(3)Hex(6)<br>HexNAc(3)Hex(5)<br>HexNAc(3)Hex(6)NeuAc(1)<br>HexNAc(3)Hex(5)NeuAc(1) | HexNAc(2)Hex(7)<br>HexNAc(2)Hex(6)<br>HexNAc(2)Hex(5) |  |
| 295-301 |  | HexNAc(2)Hex(9)<br>HexNAc(2)Hex(5) |  |
| 332-334* | HexNAc(6)Hex(6)NeuAc(1)<br>HexNAc(5)Hex(5)NeuAc(2)<br>HexNAc(6)Hex(6)NeuAc(3)<br>HexNAc(6)Hex(6)NeuAc(2) |  |  |
| 339 | HexNAc(6)Hex(6)Fuc(2)NeuAc(2)<br>HexNAc(5)Hex(6)Fuc(3)NeuAc(2) |  |  |
| 355 | HexNAc(6)Hex(4) |  |  |
| 361 | HexNAc(5)Hex(5)NeuAc(2)<br>HexNAc(6)Hex(6)NeuAc(2) |  |  |
| 448 | HexNAc(5)Hex(5)Fuc(2)NeuAc(1)<br>HexNAc(6)Hex(6)Fuc(2)NeuAc(2)<br>HexNAc(6)Hex(7)Fuc(3)NeuAc(4) | HexNAc(2)Hex(9) |  |
| 461/462 | HexNAc(5)Hex(5)<br>HexNAc(5)Hex(5)Fuc(1)<br>HexNAc(5)Hex(4)Fuc(1)<br>HexNAc(4)Hex(5)Fuc(1)NeuAc(1)<br>HexNAc(5)Hex(6)Fuc(1)NeuAc(1)<br>HexNAc(4)Hex(5)Fuc(1) |  |  |
| 616 | HexNAc(6)Hex(6)Fuc(2)NeuAc(1) |  |  |
| 637 |  |  | HexNAc(3)Hex(6) |

\* un-glycosylated peptide also observed
<sup>1</sup> Abbreviations and glycan classifications according to \*Essentials of Glycobiology\*, 3<sup>rd</sup> ed.
HexNAc: N-acetylhexosamines; Hex: Hexose; Fuc: L-Fucose; NeuAc: N-acetyl neuraminic acid

**Table S5.** Backbone carbon chain RMSD^1^ in angstroms for gp120 and gp41 subunits in high resolution Env glycoprotein structures.

| **Gp120** | 92UG037.8 full length<br>(PDB: 6ULC) | JR-FL full-length<br>(PDB: 5FUU) | AMC011 full length<br>(PDB: 6OLP) | BG505 SOSIP<br>(PDB: 4ZMJ) |
| --- | --- | --- | --- | --- |
| 92UG037.8 |  | 2.204 | 2.980 | 1.752 |
| JR-FL |  |  | 1.654 | 1.758 |
| AMC011 |  |  |  | 2.924 |
| BG505 |  |  |  |  |
| **Gp41** | 92UG037.8 | JR-FL | AMC011 | BG505 |
| 92UG037.8 |  | 5.105 | 4.888 | 3.908 |
| JR-FL |  |  | 1.928 | 2.063 |
| AMC011 |  |  |  | 1.299 |
| BG505 |  |  |  |  |

<sup>1</sup>root mean square deviation

**Table S6.** Percentage sequence similarity in Env amongst HIV-1 strains compared in Table S6.

|  | ADA-CM | Bal-1 | JR-FL | 92UG037.8 | BG505 |
| --- | --- | --- | --- | --- | --- |
| ADA-CM | 100.0 | 84.39 | 88.82 | 76.19 | 76.97 |
| Bal-1 |  | 100.0 | 89.78 | 74.68 | 77.29 |
| JR-FL |  |  | 100.0 | 76.52 | 76.04 |
| 92UG037.8 |  |  |  | 100.0 | 81.07 |
| BG505 |  |  |  |  | 100.0 |

